# ENPP1 buffers extracellular cGAMP in brown adipose tissue to limit insulin resistance

**DOI:** 10.64898/2026.04.12.718013

**Authors:** Songnan Wang, Yingjie Guo, Weidong An, Michelle Lee, Yu Li, Valentino Sudaryo, Gabriel Grenot, Gemini Skariah, Saranya C. Reghupaty, Sonny Young, Xiaochen Bai, Katrin J. Svensson, Lingyin Li

## Abstract

The ectonucleotide pyrophosphatase/phosphodiesterase 1 (ENPP1) has long been linked with metabolic diseases, with the common ENPP1 K173Q (historically K121Q) variant conferring increased risk for type 2 diabetes (T2D). However, the mechanistic basis of this association has remained unclear. Here, we demonstrate that the K173Q variant has decreased cGAMP hydrolysis activity, suggesting that this loss of enzymatic function could contribute to its pathogenesis. Using a cGAMP-hydrolysis-deficient knock-in mouse (*Enpp1^H362A^*), we show that selective loss of this activity leads to a primary defect in energy expenditure and exacerbates high-fat diet (HFD)-induced weight gain and insulin resistance. An unbiased *in vivo* glucose-uptake screen reveals brown adipose tissue (BAT) as a focal site of metabolic impairment, characterized by profound extracellular cGAMP accumulation and a selective failure of insulin-stimulated glucose uptake. Mechanistically, we demonstrate that nutrient excess drives mitochondrial DNA leakage in brown adipocytes, triggering cGAMP production and export. Excess cGAMP directly propagates STING-dependent suppression of glucose uptake and lipogenesis in brown adipocytes. Additionally, when ENPP1-mediated clearance is compromised, extracellular cGAMP acts as a paracrine immunotransmitter that remodels the BAT microenvironment by recruiting and polarizing macrophages toward an M1-like phenotype. Together, our findings nominate the impaired ENPP1-dependent buffering of extracellular cGAMP as one mechanism by which ENPP1 variants influence metabolic homeostasis.

## INTRODUCTION

Obesity and type 2 diabetes (T2D) are metabolic disorders affecting nearly 40% of the world population^1^. The increasing global prevalence has intensified the efforts to define molecular mechanisms that govern insulin sensitivity and energy homeostasis. Chronic low-grade inflammation has long been recognized as a key mediator of metabolic dysfunction^2^. For example, proinflammatory cytokines such as tumor necrosis factor (TNF-α)^3–5^ and interleukin 6 (IL-6)^6^ have been linked to insulin resistance, glucose intolerance, and T2D. However, the precise danger signals that trigger and sustain metabolic dysfunction and their regulators remain incompletely defined.

Genetic studies have long identified *ENPP1* variants, most notably the K173Q (historically K121Q) polymorphism, as significant risk factors for T2D^7–11^. For decades, the prevailing model posited that ENPP1 promotes insulin resistance by physically binding to insulin receptor (IR) at the plasma membrane and impairing its signaling transduction through a scaffolding mechanism^12^. However, this model has remained controversial due to inconsistent biochemical evidence and limited validation under physiological conditions.

An alternative explanation is that ENPP1 regulates metabolic stress-induced inflammation through its role as an immune checkpoint for extracellular 2’3’-cyclic GMP-AMP (cGAMP)^13–20^. Metabolic stress, particularly the lipotoxicity associated with a high-fat diet (HFD), causes mitochondrial damage and leakage of mitochondrial DNA (mtDNA) into the cytosol^21^. Cytosolic double stranded DNA (dsDNA) is recognized by the cyclic GMP-AMP synthase (cGAS), which catalyzes production of the second messenger cGAMP^22–25^. cGAMP then binds and activates its cytosolic sensor stimulator of interferon genes (STING), initiating a robust proinflammatory transcriptional program^22–25^. While the cGAS-STING pathway is a cornerstone of antiviral and anticancer defense^26^, recent evidence suggests it may also play a deleterious role in chronic metabolic diseases^27–29^. For example, *Sting*-deficient mice exhibit improved insulin resistance and glucose intolerance compared to wildtype (WT) mice in response to HFD^28,29^. However, most studies have focused on cell autonomous STING activation, largely leaving the fate and function of extracellular cGAMP unexplored.

Here, we show that the *ENPP1* K173Q variant has reduced cGAMP hydrolysis activity. We use a cGAMP-hydrolysis-deficient knock-in mouse (*Enpp1^H362A^*) to interrogate how ENPP1-dependent control of extracellular cGAMP influences systemic metabolism. By integrating *in vivo* metabolic phenotyping with tissue-specific glucose uptake, we identified BAT as a focal site of metabolic impairment. We demonstrate that in the absence of ENPP1 activity, extracellular cGAMP accumulates in BAT and acts as a paracrine signal that propagates STING-dependent suppression of glucose uptake and lipogenesis in brown adipocytes. Through BAT immunoprofiling, we show that ENPP1 also prevents extracellular cGAMP from activating STING signaling in macrophages and polarizing them to proinflammatory phenotypes. Together, we show that ENPP1functions as an immunometabolic checkpoint that constrains cGAMP–STING signaling in the BAT microenvironment and prevents diet-induced obesity and insulin resistance.

## RESULTS

### The common human K173Q ENPP1 allele exhibits reduced cGAMP hydrolysis activity

The K173Q ENPP1 allele has a minor homozygous allele frequency of 4.4% (**Figure 1a**). A meta-analysis of over 33,000 subjects of European descent across 20 studies found that the K173Q variant is associated with an increased risk of T2D (OR 1.38; 95% CI 1.10-1.75) under a recessive model of inheritance (**Figure 1b**)^11^. Interestingly, 16% of this genetic risk appears to be modulated by body mass index (BMI)^11^.

**Figure 1.**
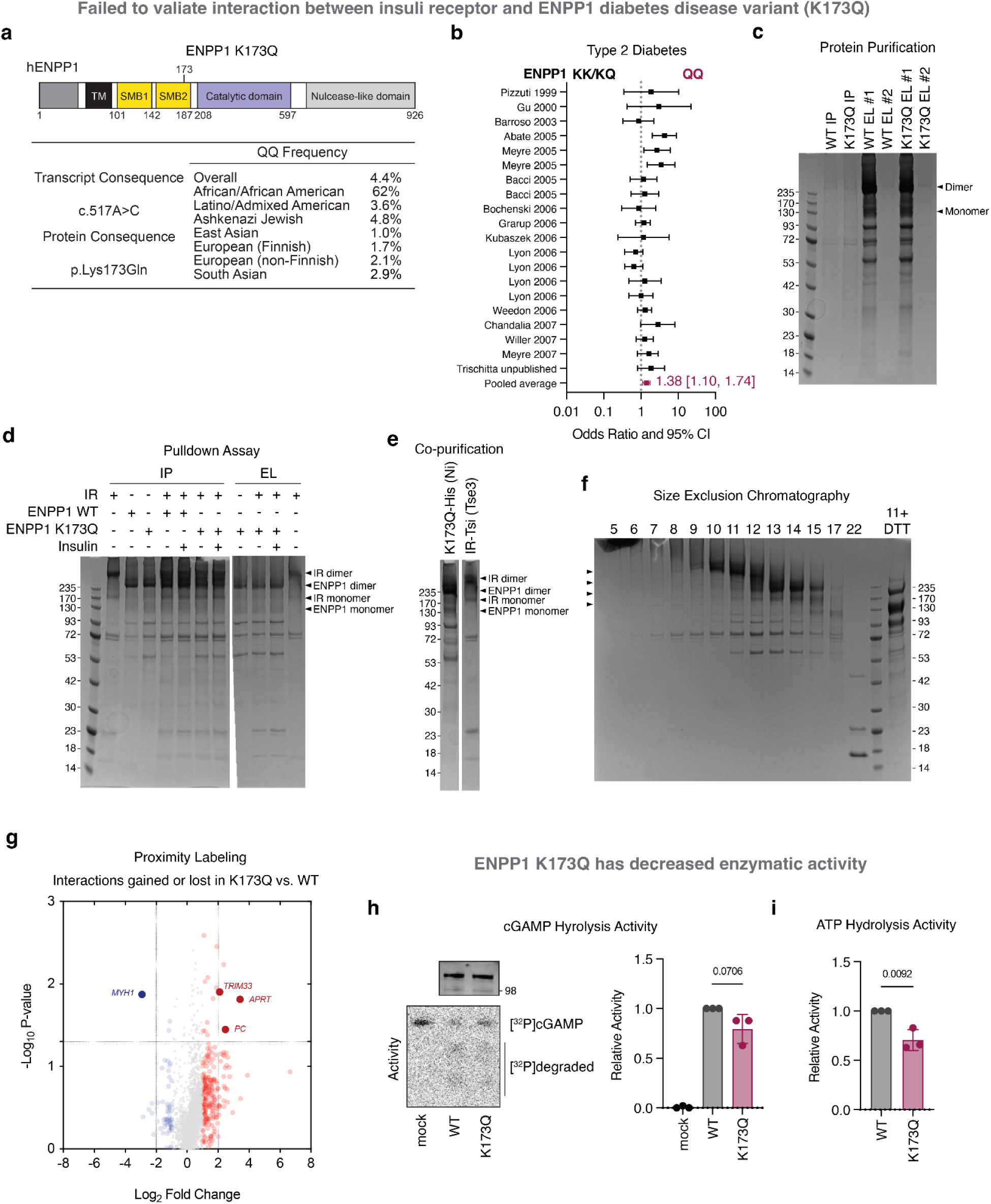
The common human K173Q ENPP1 allele exhibits reduced cGAMP hydrolysis activity. **a,** Schematic representation of the hENPP1 protein domains (top) and the allele frequency of the K173Q ENPP1 variant (bottom) **b**, Random-effects meta-analysis of 20 studies of European descent for an association of the homozygous QQ genotype with type 2 diabetes. Figure reproduced from McAteer et al^11^. **c,** Purification of His-tagged wild-type (WT) and K173Q hENPP1 from HEK293 cells via nickel-affinity chromatography. **d,** Pulldown assays of WT-hENPP1-His or K173Q-hENPP1-His with Tsi-Flag-tagged hIR (hIR-Tsi-Flag) in the presence or absence of insulin. No significant interaction was observed between hIR and the K173Q variant regardless of insulin-stimulation. **e,** Co-purification analysis of co-expressed K173Q-hENPP1-His and hIR-Tsi-Flag in HEK293 cells. No reciprocal co-purification was detected between hENPP1 and hIR. **f,** Size exclusion chromatography (SEC) of a mixture of purified K173Q-hENPP1 and hIR using a Superose 6 column. hIR (fractions 9–12) and hENPP1 (fractions 12–17) eluted as distinct peaks, indicating a lack of stable complex formation. The four arrows indicate IR dimer, ENPP1 dimer, IR monomer, and ENPP1 monomer from top to bottom. **g**, Proximity labeling using TurboID in MCF7 cells to identify the WT and K173Q hENPP1 interactomes. hIR was not detected in the proximity networks of either hENPP1 variant. Among the four proteins identified as differential interactors for the K173Q variant, hIR was not present. **h**, cGAMP degradation activity of lysates of 293T *ENPP1^-/-^* cells over expressing WT or K173Q-hENPP. Representative immunoblot and thin-layer chromatography (left) and quantification of relative enzyme activity. n = 3 independent experiments. Data are presented as mean ± s.d. **i**, Relative ATP degradation activity of WT and K173Q-hENPP1, compiled from three independent published studies^30–32^. Data are presented as mean ± s.d. hENPP1, human ENPP1; TM, transmembrane; SMB, somatomedin B domain; IR, insulin receptor; MYH1, myosin heavy chain 1; TRIM33, tripartite motif containing 33; APRT, adenine phosphoribosyltransferase; PC, pyruvate carboxylase.

Concerning the mechanism by which this variant predisposes to T2D, a prior study proposed that K173Q promotes insulin resistance by binding more strongly to IR than the WT enzyme^12^. Using multiple complementary biochemical approaches, including *in vitro* pulldowns, co-purification, size exclusion chromatography, and TurboID assays, we were unable to detect a stable interaction between K137Q ENPP1 and IR (**Figure 1c-g**). This prompted us to consider altered enzymatic activity, instead of scaffolding function, as a potential drive of the variant’s metabolic effects. Indeed, we found that human K173Q ENPP1 expressed in cell lysate has an approximately 20% reduction in cGAMP hydrolysis relative to WT (**Figure 1h**), in line with previous reports of 29% reduction of ATP hydrolysis by this variant^30–32^ (**Figure 1i**).

### Loss of ENPP1 cGAMP hydrolysis activity impairs energy balance during nutritional challenge

We then investigated whether this reduced cGAMP hydrolysis activity contributes to the increased risk associated with homozygous ENPP1 K137Q. Because this variant is not conserved in mice, and to specifically interrogate extracellular cGAMP signaling without perturbing other ENPP1 functions, we used *Enpp1^H362A^*mice, whose ENPP1 lacks cGAMP hydrolysis activity while preserving scaffolding and ATP hydrolysis^20^.

We first characterized these mice on a standard chow diet (CD) to establish a baseline metabolic profile. We measured indirect calorimetry in metabolic cages over 76 hours after one day of acclimation. Interestingly, while body weights were similar (**Figure 2a, S1a**), *Enpp1^H362A^*mice exhibited a significant reduction in oxygen consumption (VO_2_) and carbon dioxide production (VCO_2_), as determined by analysis of covariance (ANCOVA) using a general linear model (GLM) with body mass as a covariate (**Figure 2b,c**). *Enpp1^H362A^* mice also exhibited a significantly lower respiratory exchange ratio (RER = VCO_2_/VO_2_), suggesting a shift toward fatty acid oxidation from glucose utilization (**Figure 2d**). Locomotor activity was reduced in *Enpp1^H362A^* mice, whereas ambulatory movement was unaffected, indicating the absence of a primary motor deficit (**Figure 2e,f**). Consistently, mass-adjusted energy expenditure was reduced in *Enpp1^H362A^*mice, while food intake remains normal (**Figure 2g, h**). These data indicate that loss of ENPP1-mediated cGAMP hydrolysis results in a primary reduction in energy expenditure and altered substrate utilization even without nutritional stress.

**Figure 2.**
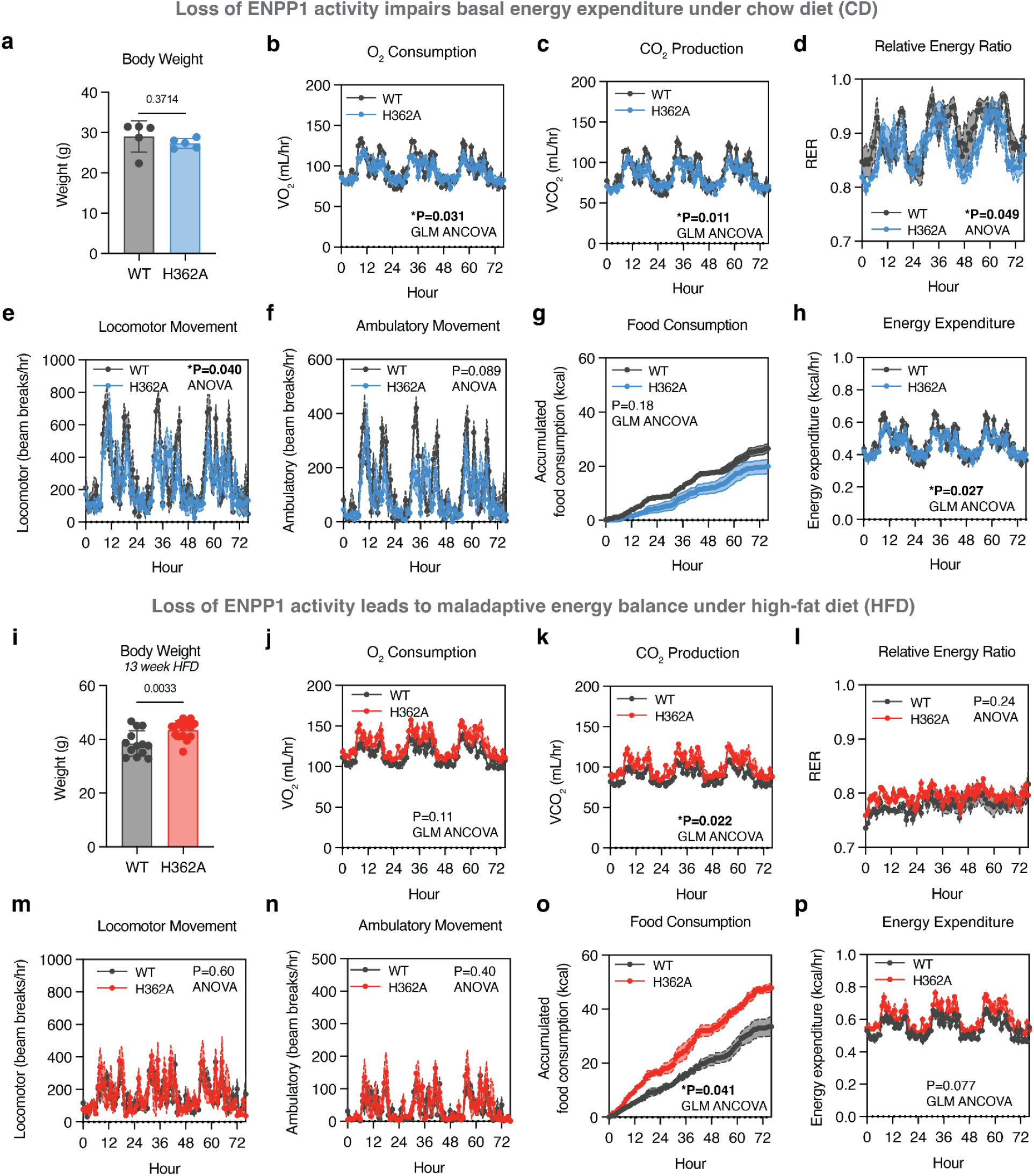
Loss of ENPP1 cGAMP hydrolysis activity impairs energy balance during nutritional challenge. **a**, Body weight of male mice on CD aged-matched to HFD cohort in **i**. Mice were housed at room temperature. *n* = 5 mice. Data are presented as mean ± s.d. **b–h**, Indirect calorimetry of hourly oxygen consumption (VO₂) (b), hourly carbon dioxide production (VCO₂) (c), hourly respiratory exchange ratio (RER) (d), hourly locomotor movement (e), hourly ambulatory movement (f), accumulated food consumption (g), and hourly energy expenditure (h), of male WT and *Enpp1^H362A^* mice housed at room temperature maintained on CD and age matched to the cohorts shown in **j-p**. Mice were acclimated for 24 h prior to data collection, and metabolic parameters were measured using a Columbus Instruments Oxymax/CLAMS indirect calorimetry system. *n* = 4 mice per group. **i**, Body weight of male mice fed a HFD for 13 weeks. Mice were housed at room temperature. *N* = 13–14 mice. Data are presented as mean ± s.d. **j–p**, Indirect calorimetry of hourly oxygen consumption (VO₂) (j), hourly carbon dioxide production (VCO₂) (k), hourly respiratory exchange ratio (RER) (l), hourly locomotor movement (m), hourly ambulatory movement (n), accumulated food consumption (o), and hourly energy expenditure (p) of male WT and *Enpp1^H362A^*mice housed at room temperature after 15 weeks of HFD feeding. Mice were acclimated for 24 h prior to data collection, and metabolic parameters were measured using a Columbus Instruments Oxymax/CLAMS indirect calorimetry system. *n* = 4 mice for WT and 3 mice for H362A (one mouse was excluded due to food consumption failed to be captured). **b-h, j-p**, Data are presented as mean ± s.m.e. Statistical significance for VO_2_, VCO_2_, food consumption, and energy expenditure was determined by General Linear Model (ANCOVA) with body mass as a covariate. Significance for RER, locomotor and ambulatory movement was determined by two-way ANOVA. See also Figure S1.

We next challenged the mice with a 60% high-fat diet (HFD) to assess their capacity to adapt energy balance to caloric excess. Although WT and *Enpp1^H362A^* exhibited similar body weights on a CD (**Figure 2a**), *Enpp1^H362A^* mice gained more weight on HFD (**Figure 2i, S1a**). We performed indirect calorimetry on male mice after 15 weeks of HFD, age-matched to the CD cohort described above. *Enpp1^H362A^* mice exhibited similar mass-adjusted VO_2_ (**Figure 2j**), higher mass-adjusted VCO_2_ (**Figure 2k**), and comparable RER to WT controls (**Figure 2l)**. No differences were observed between the genotypes in locomotor (**Figure 2m**) or ambulatory activity (**Figure 2n**). Interestingly, *Enpp1^H362A^* mice exhibited a significant increase in food consumption (**Figure 2o**), yet mass-adjusted energy expenditure was not increased (**Figure 2p**). In other words, despite higher caloric intake and greater body mass, *Enpp1^H362A^* mice failed to proportionally elevate energy expenditure, indicating impaired metabolic clearance of excess fuel, which likely exacerbated their weight gain.

This failure to scale energy expenditure was reflected in altered adipose tissue morphology. *Enpp1^H362A^* mice exhibited mildly increased inguinal white adipose tissue (iWAT) mass, while epididymal visceral adipose tissue (eVAT), interscapular BAT, and liver mass remained unchanged (**Figure S1b-e**). Histology revealed adipocyte hypertrophy in iWAT (**Figure S1f)** and more pronounced BAT hypertrophy, with larger unilocular lipid droplets in *Enpp1^H362A^*BAT (**Figure S1g**). In contrast, hepatic steatosis and non-fasting plasma free fatty acid (FFA) were similar (**Figure S1h,i**). In summary, while loss of ENPP1-mediated cGAMP hydrolysis results in a baseline deficit in energy turnover, in caloric excess, it leads to failure to proportionally scale energy expenditure with significant hyperphagia, culminating in overt obesity.

### Loss of ENPP1 cGAMP hydrolysis activity exacerbates HFD-induced insulin resistance and BAT-specific dysfunction

Given that human genetic variants in *ENPP1* are strongly associated with T2D^7–11^, we next investigated whether the loss of cGAMP hydrolysis tip systemic glucose homeostasis. Following the HFD challenge, glucose tolerance was impaired in WT mice, but *Enpp1^H362A^*mice did not show further deterioration (**Figure S2a**), and non-fasting serum glucose levels only showed a trend towards elevation (**Figure S2b**). In contrast, *Enpp1^H362A^* mice displayed dramatically worse insulin resistance compared to WT mice under HFD, as assessed by insulin tolerance tests (**Figure 3a**). Notably, while serum glucagon levels were unaffected during ad libitum feeding (**Figure S2c**), *Enpp1^H362A^* mice exhibited elevated serum insulin levels (**Figure 3b**), consistent with compensatory hyperinsulinemia to maintain near-normal glycemia despite the profound decrease in insulin sensitivity. Together, these data indicate that *Enpp1^H362A^* mice develop diet/obesity-dependent systemic insulin resistance.

**Figure 3.**
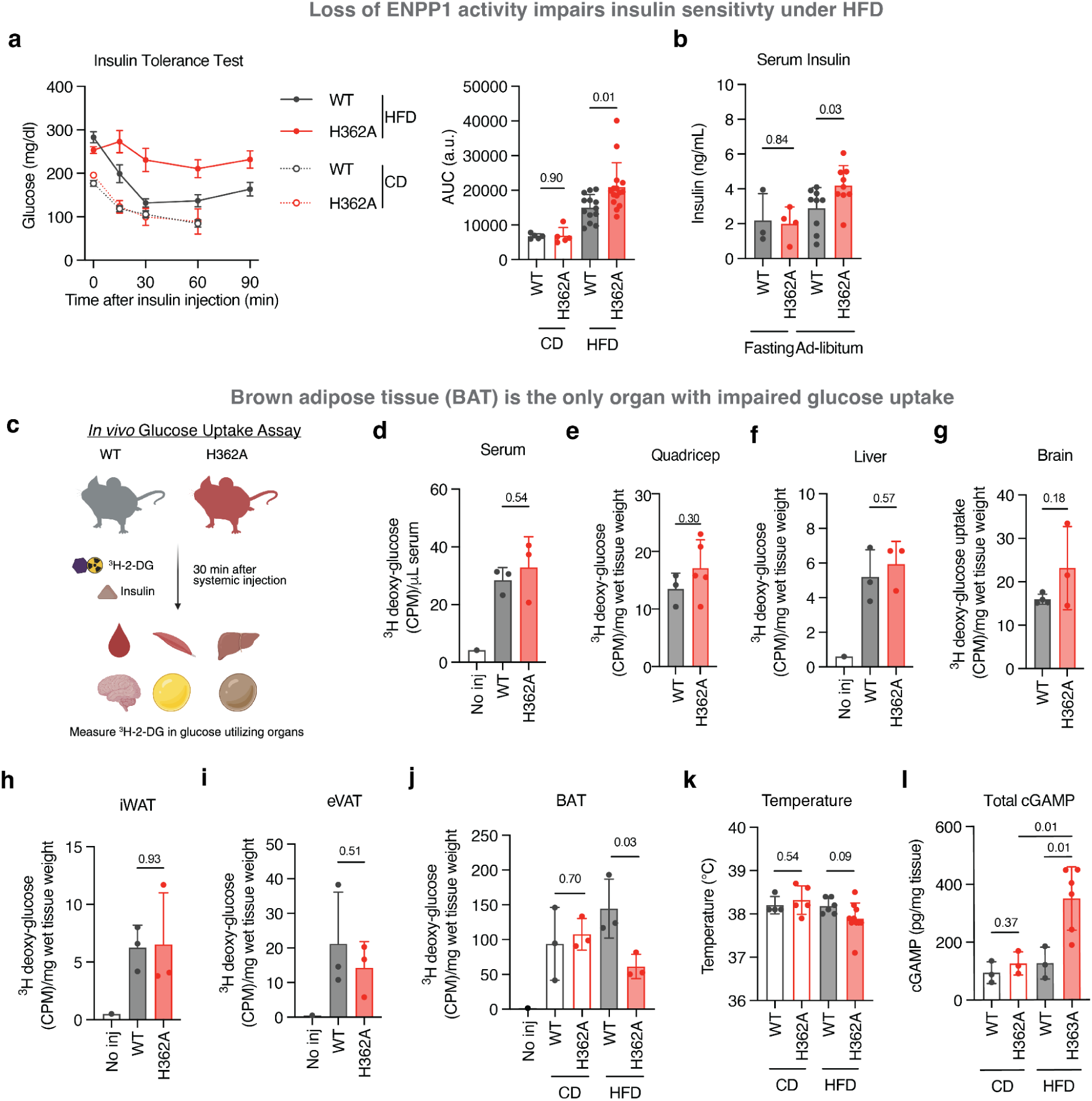
Loss of ENPP1 cGAMP hydrolysis activity exacerbates HFD-induced insulin resistance and BAT-specific dysfunction. **a**, Insulin tolerance tests (ITTs) performed after 13 weeks of HFD or CD feeding in male mice housed at room temperature. Mice were fasted for 5–6 h prior to intraperitoneal injection of insulin (1 U kg⁻¹). Blood glucose concentrations were measured over time and AUC was calculated. *n* = 5 mice for CD and *n* = 13–15 mice for HFD. Line graphs show mean ± s.e.m.; bar graphs show mean ± s.d. **b**, Serum insulin concentrations in male mice housed at room temperature after 8, 12 or 14 weeks of HFD feeding under fasted or ad libitum conditions. Data are presented as mean ± s.d. **c**, Schematic of *in vivo* glucose uptake assay. **d-j,** *In vivo* ^3^H-2-DG uptake in serum, quadricep, liver, brain, iWAT, eVAT, and BAT of male WT and H362A mice housed at room temperature fed with HFD for 14 weeks (and their age-matched CD controls in BAT). Mice were fasted for 6 h and injected with 1 U kg⁻¹ insulin 30 m before ^3^H-2-DG measurement. Mice whose blood glucose dropped <50% from baseline were excluded.*n* = 3 mice per group except for quadriceps from H362A mice (*n* = 5). Data are presented as mean ± s.d. **k,** Rectal temperature of male WT and H362A mice housed at room temperature on 20 weeks of HFD or age-matched CD control. *n* = 4-10 mice per group. Data are presented as mean ± s.d. **i,** Total cGAMP levels in BAT from male mice on 15 weeks of HFD or age-matched CD control. Tissues were homogenized in 100 mg mL⁻¹ T-PER lysis buffer with ENPP1 inhibitor (15 µM STF-1084), diluted 100-fold, and measured by ELISA. *n* = 3-6 mice per group. Data are presented as mean ± s.d. Statistical significance was assessed using two-sided unpaired *t* tests unless otherwise indicated. CPM: counts per minute. See also Fig. S2.

To pinpoint the tissues responsible for this defect, we conducted an *in vivo* glucose uptake assay using 2-deoxy-D-[1,2-^3^H]glucose (^3^H-2-DG)^33^ (**Figure 3c**). Residual serum radioactivity was comparable between genotypes (**Figure 3d**), indicating similar tracer availability. Tissue analysis revealed that insulin-stimulated glucose uptake in skeletal muscle, liver, brain, iWAT, and eVAT did not differ between WT and *Enpp1^H362A^* mice (**Figure 3d-3i**). In contrast, BAT from *Enpp1^H362A^* mice showed a significant reduction in insulin-stimulated ^3^H-2-DG uptake following HFD (**Figure 3j**). Because BAT is a major glucose sink supporting non-shivering thermogenesis at temperatures at sub-thermoneutral temperatures, we measured core body temperature at room temperature (20-25°C). We observed a trend toward cooler rectal temperatures in HFD-fed *Enpp1^H362A^* mice (**Figure 3k**), consistent with impaired thermogenic function of the BAT.

Finally, to directly link these defects to cGAMP handling, we quantified tissue cGAMP levels. Under CD, cGAMP concentrations were similar between genotypes. After HFD, however, BAT from *Enpp1^H362A^* mice had a profound accumulation of total cGAMP levels (∼500 nM) compared to WT counterparts (**Figure 3i**). Together, these results indicate that ENPP1-mediated clearance of extracellular cGAMP is essential for maintaining insulin-stimulated glucose uptake and thermogenetic capacity in BAT during nutritional stress. Based on these findings, we focused our subsequent investigation on mechanisms by which the ENPP1-cGAMP axis maintains the BAT microenvironment and protects against tissue-wide insulin resistance.

### Brown adipocytes are a source of extracellular cGAMP driven by excess cytosolic mtDNA

Metabolic stress, such as HFD-induced lipotoxicity, is associated with mitochondrial dysfunction and the subsequent leakage of mtDNA into the cytosol^34,35^. Given the high mitochondrial density characteristic of brown adipocytes^36^, we hypothesized that they also harbor high cytosolic mtDNA. To test this, we isolated the stromal vascular fraction (SVF) of BAT from WT mice and differentiated the preadipocytes into mature brown adipocytes, confirmed by the induction of uncoupled protein 1 (*Ucp1*) (**Figure 4a**). We performed subcellular fractionation to isolate the cytosolic fractions. The purity of this fraction was confirmed by enrichment of the cytosolic marker glyceraldehyde-3-phosphate dehydrogenase (GAPDH) and absence of the mitochondrial outer membrane protein TOM20 and the nuclear marker histone 3 (H3) (**Figure 4b**). Quantitative PCR (qPCR) of the cytosolic fraction revealed approximately 10-fold higher cytosolic mtDNA levels in brown adipocytes than in preadipocytes across multiple mitochondrial loci, including cytochrome b (Cytb), D-loop, mitochondrial tRNA phenylalanine (mt-Tf), and NADH dehydrogenase 4 (ND4) (**Figure 4b**).

**Figure 4.**
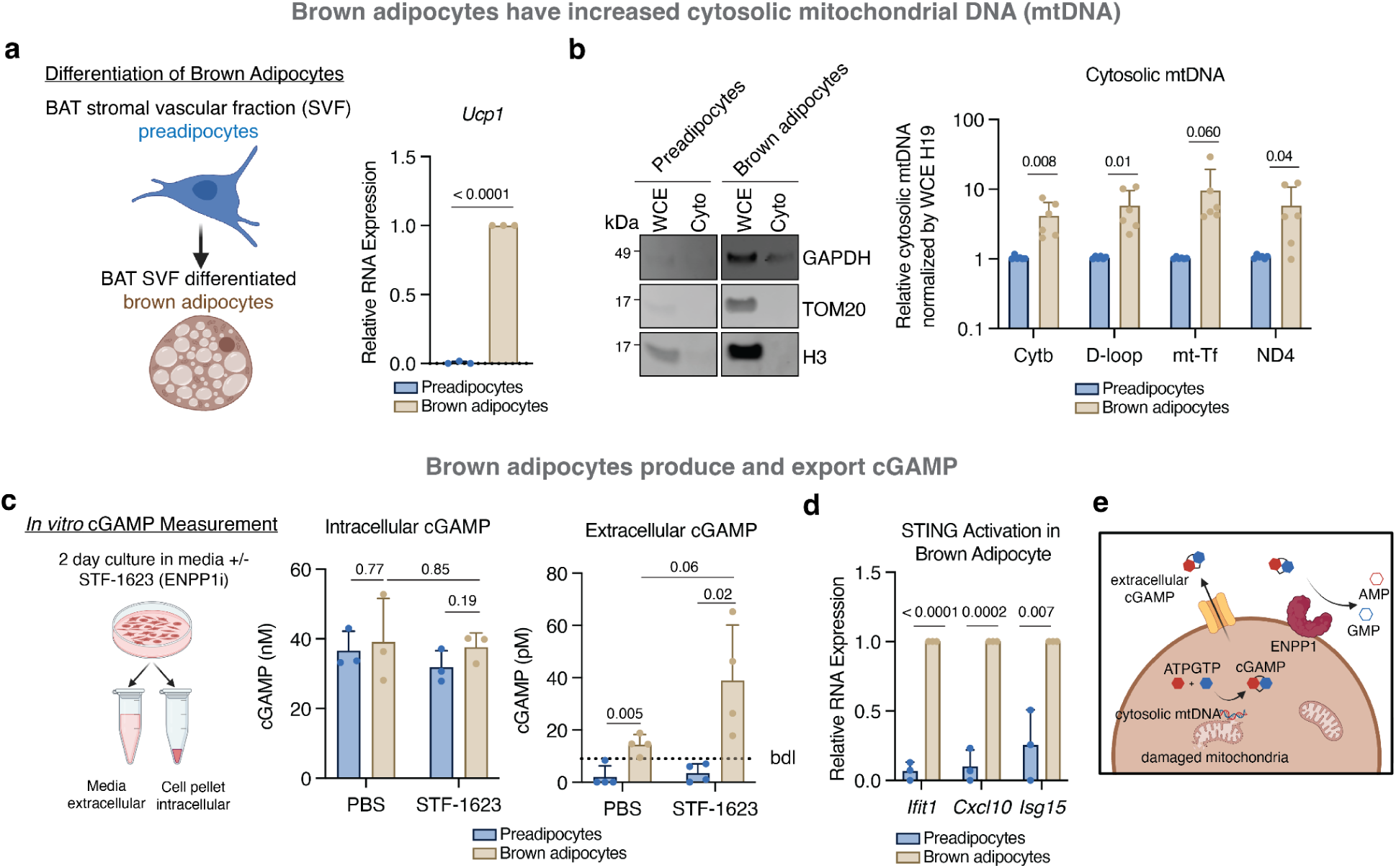
Brown adipocytes are a source of extracellular cGAMP driven by cytosolic mtDNA. **a**, Schematic of brown adipocyte differentiation (left), and relative *Ucp1* expression as a marker of brown adipocytes in primary brown preadipocytes and differentiated brown adipocytes, normalized to levels in differentiated brown adipocytes (right). *n* = 3 independent experiments. Data are presented as mean ± s.d. **b,** Western blot of WCE and cytosolic fractions from brown preadipocytes and differentiated brown adipocytes using cytosolic marker GAPDH, mitochondrial marker TOM20 and nuclear marker H3 (left). Relative cytosolic mtDNA markers (Cytb, D-loop, mt-Tf, ND4) normalized to H19 in WCE, then normalized to levels of brown preadipocytes (right). *n* = 2 independent experiments, 3 biological replicates each. Data are presented as mean ± s.d. **c,** Schematic of *in vitro* cGAMP measurement (left). Intracellular cGAMP in lysates of brown preadipocytes and differentiated brown adipocytes cultured with or without ENPP1 inhibitor (1 µM STF-1623) over 2 days (middle). Extracellular cGAMP in supernatants of brown preadipocytes and differentiated brown adipocytes cultured with or without ENPP1 inhibitor (1 µM STF-1623) over 2 days (right). bdl, 9 pM. *n* = 3 (intracellular) or 4 (extracellular) independent experiments. Data are presented as mean ± s.d. **d,** Relative *Ifit1*, *Cxcl10*, and *Isg15* expression as markers of cGAMP-STING pathway activation in primary brown preadipocytes and differentiated brown adipocytes, normalized to levels in differentiated brown adipocytes. *n* = 3 independent experiments. Data are presented as mean ± s.d. **e,** Schematic model of brown adipocytes as a source of extracellular cGAMP driven by cytosolic mtDNA. Statistical significance was assessed using two-sided unpaired *t* tests. WCE, whole cell extract; cyto, cytosolic.

We then quantified intra- and extracellular cGAMP levels in preadipocytes and brown adipocytes, with and without the ENPP1 inhibitor STF-1623^37^ (**Figure 4c**). While intracellular cGAMP levels remained stable, extracellular cGAMP concentrations were >10-fold higher in brown adipocytes than in preadipocytes (38.9 pM vs. 3.5 pM; **Figure 4c**). Notably, extracellular cGAMP was detectable even when subject to degradation by endogenous ENPP1 from these cells and 10% serum in cell culture media, and its levels increased by 2.5-fold with STF-1623 (**Figure 4c**), mirroring the effect of *Enpp1^H362A^*on cGAMP concentrations in BAT (**Figure 3i**) . While bulk extracellular cGAMP measurements in the cell culture system reflect an arbitrary 1000-fold medium dilution, the *in vivo* pericellular cGAMP concentration is likely substantially higher, as evidenced by the cGAMP accumulation measured in BAT (**Figure 3i**). Furthermore, while these *in vitro* assays capture a limited period of extracellular cGAMP accumulation, *Enpp1^H362A^*mice experience chronic, constitutive exposure. Consistent with functional signaling, brown adipocytes displayed significantly higher expression of interferon-stimulated genes (ISGs), including interferon-induced protein with tetratricopeptide repeats 1 (*Ifit1*), interferon gamma-induced protein 10 (*Cxcl10*), and interferon-stimulated gene 15 (*Isg15*) (**Figure 4d**). Together, these data indicate that microchondria-rich brown adipocytes have an elevated cytosolic mtDNA load and constitutively produce physiologically meaningful amounts of cGAMP, establishing them as a source of tonic extracellular cGAMP-STING signaling in BAT (**Figure 4e**).

### cGAMP-STING signaling induces insulin resistance and impairs glucose uptake in brown adipocytes

We next asked whether elevated cGAMP seen in *Enpp1^H362A^* BAT under HFD impairs brown adipocyte glucose handling. First, we tested whether *Enpp1^H362A^* brown adipocytes have an intrinsic defect in glucose uptake when cultured in complete media with 10% serum, where extracellular cGAMP is subject to degradation by serum ENPP1. Under both basal and insulin-stimulated conditions, *Enpp1^H362A^* and WT brown adipocytes showed comparable glucose uptake (**Figure S3a**), indicating that ENPP1 activity loss does not create a cell-intrinsic uptake defect.

We then directly examined how cGAMP-STING signaling affects brown adipocyte glucose uptake (**Figure 5a**). cGAMP treatment marked reduced glucose uptake in WT brown adipocytes under both basal and insulin-stimulated conditions (**Figure 5b,c**). This effect was completely absent in *Sting^-/-^* brown adipocytes, indicating that the suppression of glucose uptake is mediated by STING activation rather than byproducts of cGAMP hydrolysis, such as adenosine (**Figure 5b,c**). The suppression was dose-dependent, detectable at concentrations as low as 500 nM (**Figure 4d,e**), and occurs without measurable loss of cell viability (**Figure 5e**).

**Figure 5.**
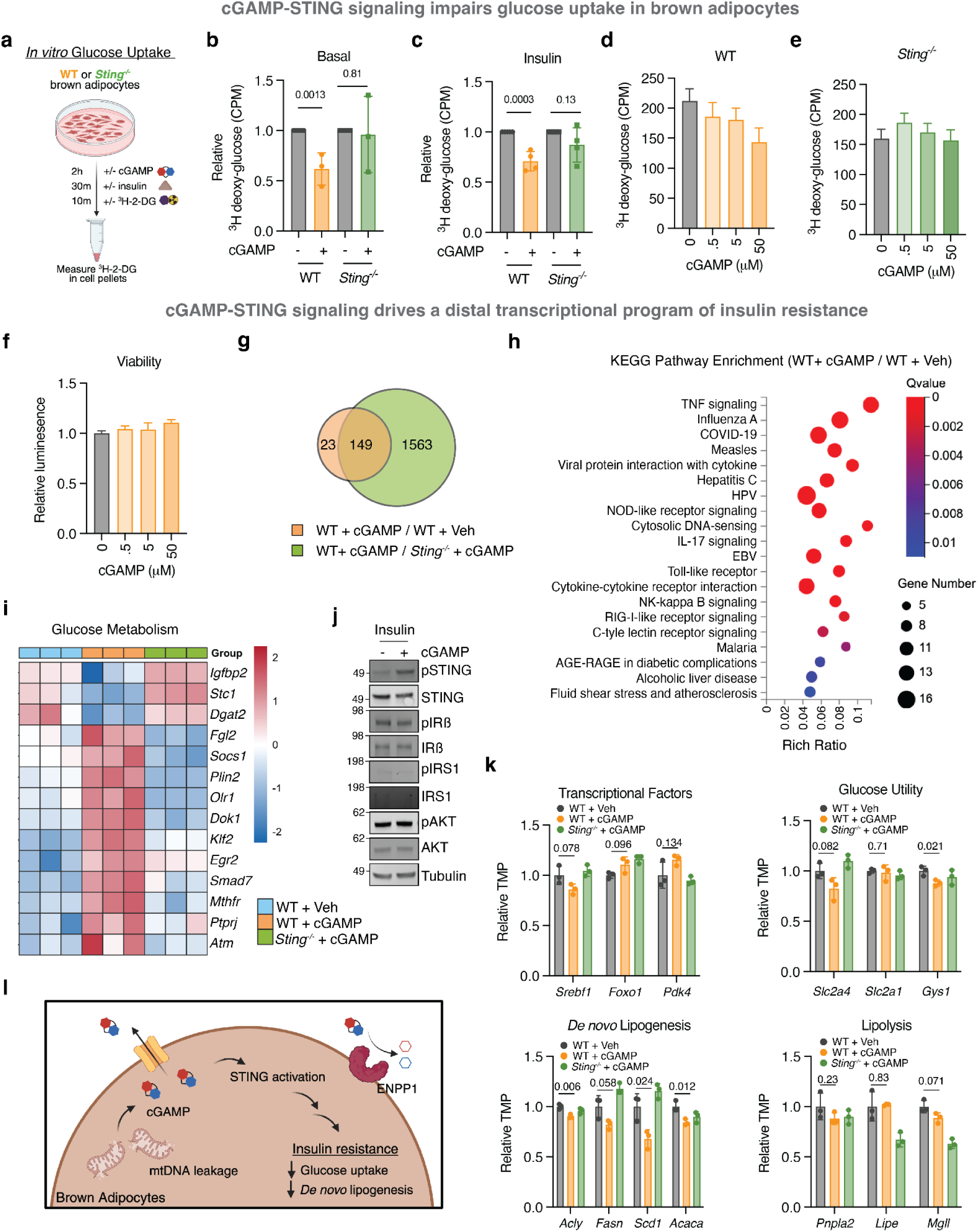
cGAMP-STING signaling induces insulin resistance and impairs glucose uptake in brown adipocytes. **a,** Schematic of *in vitro* glucose uptake assay. **b–c,** Normalized ^3^H-2-DG uptake in WT and *Sting^-/-^*brown adipocytes treated with 50 µM cGAMP for 2 h under basal conditions (**b**) or following 20 min insulin (100 nM) treatment (**c**). Data normalized to untreated control. *n* = 3 or 4 independent experiments. Data are presented as mean ± s.d. **d–e,** ^3^H-2-DG uptake in WT (**c**) or *Sting^-/-^*(**d**) brown adipocytes treated with 50–50,000 nM cGAMP for 2 h. *n* = 5 biological duplicates from 3 independent experiments. Data are presented as mean ± s.d. **f,** Cell viability of WT brown adipocytes treated with 50–50,000 nM cGAMP for 2 h, measured by CellTiter-Glo. *n* = 4 biological replicates from 2 independent experiments. Data are presented as mean ± s.d. **g-i, k**, Bulk RNA sequencing of WT brown adipocytes treated with vehicle (WT + Veh), WT brown adipocytes treated with cGAMP (WT + cGAMP), and *Sting^-/-^* brown adipocytes treated with cGAMP (*Sting^-/-^* + cGAMP). Cells were treated with 50 µM cGAMP for 2 h and, where indicated, and 100 nM insulin for 30 min. *n* = 3 biological replicates per group. Sequencing was performed as pair-ended reads on a DNBseq-G400 platform. Expression was normalized to transcripts per million (TPM). Significantly altered genes were defined as Q < 0.05. **g**, Venn diagram of significantly altered genes between WT + Veh vs. WT + cGAMP and WT + cGAMP vs. *Sting^-/-^* + cGAMP. **h**, GO-KEGG pathway enrichment analysis of genes significantly altered between WT + Veh vs. WT + cGAMP. Color scale represents Q-value. **i**, Heat map of significantly altered genes involved in glucose metabolism. Color scale represents z-scores. **j**, Western blot of WT brown adipocytes treated with or without 50 µM cGAMP for 2 h, and with 100 nM insulin for 30 min. cGAMP activation was assessed by pSTING, and insulin signaling by pIRβ, pIRS1, and pAKT expression. **k**, Relative TMP of genes regulated by insulin signaling, including transcriptional factors, genes involved in glucose utilization, lipogenesis, and lipolysis. *n* = 3 biological replicates per group. Data are presented as mean ± s.d. **l**, Schematic model of cGAMP-STING signaling inducing insulin resistance and impairing glucose uptake in brown adipocytes. Statistical significance was assessed using two-sided unpaired *t* tests unless otherwise noted. GO-KEGG: Gene Ontology-Kyoto Encyclopedia of Genes and Genomes; TMP,: transcripts per million See also Fig. S3.

To define the transcriptional basis for STING-signaling’s effect on insulin-dependent glucose uptake, we performed bulk RNA sequencing (RNAseq) on WT brown adipocytes treated with vehicle or cGAMP, using *Sting^-/-^*brown adipocytes treated with cGAMP as a control. Almost all cGAMP-induced changes in gene expression were STING-dependent (**Figure 5g**). Pathways analysis revealed that cGAMP primarily perturbed immune and cardiometabolic pathways, including AGE-RAGE signaling^38^, alcoholic liver disease, and fluid shear stress and atherosclerosis pathways (**Figure 5h)**. As expected, cGAMP robustly induced ISGs (**Figure S3b**). Notably, even at baseline WT brown adipocytes exhibited higher ISG expression than *Sting^-/-^*cells (**Figure S3c**), consistent with tonic endogenous cGAMP production from basal cytosolic mtDNA (**Figure 5b,c**). In contrast, pathways such as TNF, NF-κB, and AGE-RAGE were only induced by exogenous cGAMP, suggesting they require a higher activation threshold than basal ISG signaling (**Figure S3c**). Importantly, cGAMP also altered key regulators of glucose metabolism in a STING-dependent manner (**Figure 5i**): it downregulated stanniocalcin 1 (*Stc1*), a positive regulator of glucose uptake in BAT^39^, and upregulated perilipin 2 (*Plin2*), which can impair glucose uptake by disrupting the SNARE machinery required for GLUT4 translocation^40^.

We next examined where STING signaling intersects with the insulin signaling cascade. In primary brown adipocytes, cGAMP treatment did not affect phosphorylation of the insulin receptor β-subunit (pIRβ), insulin receptor substrate 1 (pIRS1), or the downstream kinase AKT (pAKT), indicating that the proximal signaling remains functional (**Figure 5f**). In contrast, transcriptomic profiling revealed a marked impairment of the distal insulin-regulated transcriptional network. cGAMP-STING signaling significantly blunted the activity of the AKT-mTORC1-SREBP1c axis, as evidenced by the downregulation of the master regulator sterol regulatory element binding transcription factor 1 (*Srebf1*) and its lipogenic targets, including ATP citrate lyase (*Acly),* fatty acid synthase *(Fasn*), stearoyl-CoA desaturase 1 (*Scd1*), and acetyl-CoA carboxylase alpha (*Acaca* or *Acc1*)^41^ (**Figure 5k**). Furthermore, insulin-mediated suppression of forkhead box O1 (*FoxO1*) and target genes appeared compromised, as shown by elevated levels of *FoxO1* and its target pyruvate dehydrogenase kinase 4 (Pdk4)^42^ (**Figure 5k**). Consistent with impaired insulin-driven glucose utilization, expression of the insulin-responsive glucose transporter GLUT4 (*Slc2a4*) and glycogen synthase 1 (*Gys1*) was reduced, whereas the basal glucose transporter GLUT1 (*Slc2a1*) was unchanged (**Figure 5k**). Finally, the anti-lipolytic arm of insulin action was comparable between groups, including adipose triglyceride lipase (*Pnpla2* or *Atgl*), hormone-sensitive lipase (*Lipe*), and monoglyceride lipase (*Mgll*) (**Figure 5k**). Taken together, these data are consistent with a model in which cGAMP-STING signaling does not block proximal insulin signaling, but instead reprograms the distal transcriptional response: STING activation suppresses the AKT-mTORC1-SREBP1c axis and induces *FoxO1*-*Pdk4* expression, while modulating regulators of glucose uptake, such as *Stc1* and *Plin2.* Together, these changes shift brown adipocytes into a transcriptional state of insulin resistance, characterized by the inhibition of anabolic programs governing glucose uptake and lipogenesis despite preserved AKT phosphorylation. In the setting of impaired ENPP1-mediated cGAMP clearance, HFD-stressed brown adipocytes exhibit STING-dependent metabolic defects to initiate a vicious cycle (**Figure 5l**).

### Paracrine cGAMP-STING signaling recruits and polarizes M1 macrophages

Beyond direct effects on brown adipocytes, we next investigated whether other cell types within the BAT microenvironment are affected by the accumulation of extracellular cGAMP. Because cGAMP can act as an immunotransmitter^14^, we focused on immune composition in BAT from HFD-fed *Enpp1^H362A^* mice, using iWAT as an adipose control. Flow cytometry analysis revealed a significant increase in percentage of total CD45^+^ immune cells specifically in the BAT, but not in the iWAT, compared to WT controls (**Figure 6a,b**). This expansion was largely accounted for by a higher percentage of macrophages (**Figure 6c,d**), whereas the frequencies of monocytes, neutrophils, dendritic cells, T cells, B cells, and NK cells were similar between genotypes in both BAT and iWAT (**Figure S4**). These results are consistent with the established paradigm of chronic overnutrition leading to macrophage infiltration^43,44^, but also point to a BAT-selective, extracellular cGAMP role in immunemetabolic remodeling..

**Figure 6.**
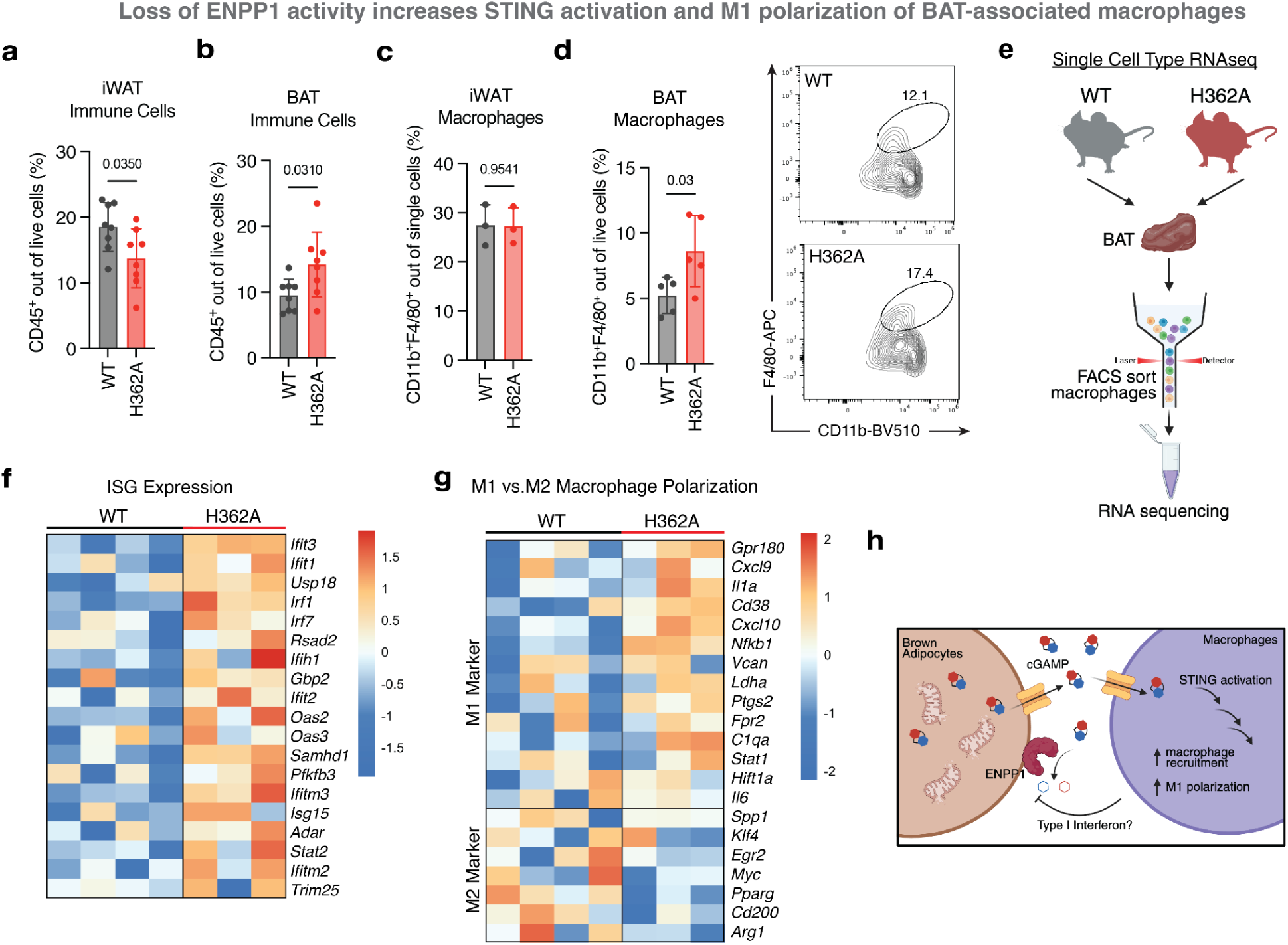
Paracrine cGAMP-STING signaling recruits and polarizes M1 macrophages. **a** and **b**, Percentage of immune cells (CD45⁺) among live cells in the iWAT (**a**) and BAT (**b**) of male WT and H362A mice housed at room temperature and fed a HFD for 16 weeks. *n* = 4 mice per group; two technical replicates per mouse (left and right depots). Data are presented as mean ± s.d. **c**, Percentage of macrophages (CD11b⁺F4/80⁺) among live cells in iWAT from male WT and H362A mice housed at room temperature and fed an HFD for 15 weeks. *n* = 3 mice per group. Data are presented as mean ± s.d. **d**, Left, representative flow cytometry plots showing macrophages (CD11b⁺F4/80⁺) gated within CD11b⁺ cells in BAT from male WT and H362A mice housed at room temperature and fed an HFD for 9 weeks. Right, percentage of macrophages among live cells. *n* = 5 mice per group. Data are presented as mean ± s.d. **e**, Schematic of RNA sequencing of BAT associated macrophages. **f** and **g**, Bulk RNA sequencing of 1,000 FACS-sorted BAT-associated macrophages (CD11b⁺F4/80⁺) isolated as in **(e)** WT and H362A mice. Only samples passing quality control after library preparation were sequenced. Expression of ISGs (**f**) and M1- and M2-associated macrophage markers (**g**). Heat maps are scaled by z-score. **g**, Schematic model of paracrine cGAMP-STING signaling recruiting and polarizing M1 macrophages. Statistical significance was assessed using two-sided unpaired ***t*** tests unless otherwise noted. See also Fig. S4.

Macrophage polarization state is a critical determinant of brown adipocyte function: while M2 (anti-inflammatory, alternatively activated) macrophages typically promote BAT thermogenesis, M1 (pro-inflammatory, classically activated) macrophages are known to impair it^45–48^. To assess the gene expression profile and polarization state of these cells *in vivo*, we sorted BAT-associated macrophages from WT and *Enpp1^H362A^*mice and performed RNA sequencing (**Figure 5e**). We observed the elevated expression of ISGs in BAT-associated macrophages from *Enpp1^H362A^* mice (**Figure 5e**), consistent with a STING-mediated response to increased extracellular cGAMP. Furthermore, transcriptomic analysis revealed a distinct phenotypic shift from an M2-like toward an M1-like state in macrophages from the *Enpp1^H362A^* mice relative to WT controls (**Figure 5f**), providing a further mechanistic basis for the BAT dysfunction observed in these mice. Interestingly, we previously observed that extracellular cGAMP in the tumor microenvironment drives a similar M2 to M1 repolarization of tumor-associated macrophages, suggesting a conserved role for extracellular cGAMP in promoting pro-inflammatory macrophage states across diverse physiological contexts^49^.

Together with our brown adipocyte-intrinsic data, we propose a model in which mitochondria-rich brown adipocytes produce and export cGAMP in response to cytosolic mtDNA leakage under conditions of nutrient excess (**Figure 4e**). Under normal conditions, ENPP1 continuously degrades extracellular cGAMP, limiting its concentrations, spatial range and duration. When ENPP1 activity is compromised, cGAMP accumulates in BAT and drives a two-pronged failure of the tissue: it directly induces a distal insulin-signaling defect and impairs glucose uptake in brown adipocytes (**Figure 5l**), and it simultaneously recruits and polarizes macrophages toward a M1-like pro-inflammatory state; type I interferon response in these cells cause further mitochondria damage and impairment in brown adipocyte function, feeding into the vicious cycle^50^ (**Figure 6h**). These convergent adipocyte-intrinsic and immune-mediated mechanisms together compromise BAT metabolic function. In this framework, ENPP1 functions as an immunometabolic checkpoint that preserves BAT function and, by extension, contributes to maintenance of systemic energy balance.

## DISCUSSION

A key observation of our study is the double hit requirement for over metabolic disease. Although *Enpp1^H362A^* mice exhibit a baseline reduction in energy expenditure on CD, classical features of metabolic dysfunction, including obesity and insulin resistance, emerge only under the stress of a HFD. This divergence is explained by a diet-dependent accumulation of cGAMP. On CD, extracellular cGAMP remains low, whereas HFD triggers a substantial increase in cGAMP production and export from brown adipocytes that cannot be buffered in the absence of ENPP1 activity, leading to impaired BAT metabolic activity. These findings support a model in which ENPP1 acts as a high-capacity extracellular cGAMP buffer that is particularly important for preserving BAT function during periods of metabolic stress. Once considered relevant primarily in infants, BAT is now recognized as a major regulator of adult human metabolism, with studies showing marked increase in serum glucose-clearing capacity under cold exposure^51,52^ and inverse correlation between BAT mass and body mass index (BMI)^53^, fasting glucose, and hemoglobin A1c levels^54^.

While our work focuses on ENPP1-mediated regulation of cGAMP in energy expenditure, the systemic obesity observed in *Enpp1^H362A^* mice on HFD reflects a combination of reduced energy expenditure and marked hyperphagia. The loss of appetite control suggests that the ENPP1-cGAMP axis may also influence central energy regulation. Recent studies have implicated the cGAS-STING pathway in age-related inflammation and neurodegeneration^55^, raising the possibility that excessive extracellular cGAMP-STING signaling in the hypothalamus could contribute to leptin resistance and disordered hunger and satiety signaling. Although this remains speculative and was not directly tested here, our data are consistent with a broader view in which ENPP1 helps limit f extracellular cGAMP-driven inflammation across multiple organ systems, extending the concept of cGAMP paracrine signaling beyond anti-cancer and anti-viral immunity^20,49^.

For decades, the association between the *ENPP1* K173Q variant and T2D has been controversial^7–11^. The prevailing model proposed that ENPP1, insteading of functioning as an enzyme, promotes insulin resistance by directly binding the IR and inhibiting its signaling through a scaffolding mechanism^12^. In contrast, our biochemical analysis did not support a stable ENPP1-IR interaction. Instead, we find that the K173Q variant exhibits reduced cGAMP hydrolysis activity. This suggests an alternative, non-mutually exclusive explanation: at least part of the diabetes risk associated with the variation may arise from impaired clearance of extracellular cGAMP, analogous to the mechanism revealed in the *Enpp1^H362A^* mouse model. We emphasize that this likely represents one contributing mechanism rather than a complete account of ENPP1 genetics in human T2D.

More broadly, our findings add to growing evidence that innate immune pathways and metabolic regulation are tightly intertwined. ENPP1 is emerging as an important innate immune checkpoint of the cGAS-STING axis^49,56^ and our work positions this checkpoint within an immunometabolic framework in BAT. Modulating ENPP1 or the cGAS-STING signaling pathway therefore has potential as a therapeutic strategy for mitigating metabolic dysfunction, but will require clever targeting strategies given the central roles of these players in anti-cancer and anti-viral immunity. Ongoing efforts to develop ENPP1-based biologics^57^, as well as inhibitors of cGAS ^55,58,59^ and STING^60–62^ underscore the clinical interest in this signaling axis. Beyond metabolic disorders, unwanted chronic or acute cGAS-cGAMP-STING activation has been implicated in aging and neurodegeneration^55^, myocardial infarction^63,64^, and various autoimmune syndromes^65^. Future studies should test whether ENPP1 similarly contributes to immune homeostasis in these contexts and to what extent impaired extracellular cGAMP clearance represents a shared mechanism across inflammatory and metabolic diseases.

## Methods

### Mouse strains

C57BL/6J (#000664) and C57BL/6J-Sting1gt/J (#017537, denoted as *Sting^-/-^*) mice were purchased from the Jackson Laboratory and B6.*Enpp1^H362A^* mice were generated in a previous publication^20^. Male mice were used for all experiments. For high fat diet (HFD) groups, mice between 4-6 weeks of age were fed with 60 kcal% fat diet (Research Diets, #D12492). Mice maintained at Stanford University at room temperature (20-25°C) followed the Stanford University Institutional Animal Care and Use Committee (IACUC) regulations. All procedures done at Stanford were approved by the Stanford University Administrative Panel on Laboratory Animal Care (APLAC Protocol No. 30591).

### Brown adipocyte culture

Brown preadiapocyte isolation and brown adipocyte differentiation were adapted from previous publications^66–69^. Briefly, interscapular brown adipose tissue (BAT) from five 4-week-old WT, *Enpp1^H362A^*, or *Sting^-/-^* male mice were pooled and incubated in 5 mL digestion buffer consisted of 0.1% collagenase type 2 (Sigma-Aldrich, C6885) in DMEM and 1% penicillin-streptomycin at 37°C in a shaking water bath for one hour. The solution was cooled at 4 °C for 20 minutes. Infranatant was filtered through a 100 µM filter and centrifuged for 5 minutes at 700 xg. The digestion buffer was removed, and the pellet was washed with DMEM and 1% penicillin-streptomycin before centrifuging for 5 minutes at 700 xg. Pellet was resuspended in 0.6 mL DMEM and 1% penicillin-streptomycin. 0.2 mL of suspension was added in each 6-well with 1.8 mL DMEM and 1% penicillin-streptomycin media. On the next day, cells were immortalized through retroviral expression of SV50 Large T-antigen (Addgen, #18922). Immortalized brown preadipocytes were selected with 0.5-1 µg puromycin between day 5-12 after cell isolation. Brown preadipocytes are passaged in maintenance media consisting of 20 nM insulin (Sigma-Aldrich, I1882), 1 nM T3 (Sigma-Aldrich, T6367), and 10% FBS in DMEM. For differentiation into mature brown adipocytes, brown preadipocytes were seeded in maintenance media on day 0 aiming for 80-90% confluence on day 1, differentiation induction media consisted of 20 nM insulin, 1nM T3, 0.5mM 3-isobutyl-1-methylxanthine (IBMX, Sigma, #410857), 0.5 µM dexamethasone (Sigma-Aldrich, D4902), 125 µM indomethacin (Sigma-Aldrich, I7378), 1µM rosiglitazone (Sigma-Aldrich, R2408), and 10% FBS in DMEM was replaced on day 2. On day 4, differentiation media consisted of 20 nM insulin, 1 nM T3, 1 µM rosiglitazone, and 10% FBS in DMEM was replaced. Cells are ready for experiment on day 6. Successful differentiation into brown adipocytes was confirmed with cell morphology (multilocular cytoplasm) and *Ucp1* expression by qRT-PCR (see notes below). All cells were maintained in a humidified incubator at 37°C and 5% CO_2_. All cell lines tested negative for mycoplasma contamination.

### Glucose tolerance and insulin tolerance test (GTT and ITT)

Male mice were fasted for 5-6 hours beginning in the early light phase with free access to water. Baseline blood glucose levels were measured from tail vein blood using a handheld glucometer (ReliOn Premier). For glucose tolerance tests, D-glucose (Sigma-Aldrich, #50-99-7) was dissolved in sterile saline and administered at a dose of 1.5 g kg^-1^ body weight by intraperitoneal injection. Blood glucose concentrations were measured at 15, 30, 60, 90, and 120 min after glucose administration. For insulin tolerance tests (ITT), human insulin (Humulin R, Eli Lilly) was diluted in sterile saline and administered intraperitoneally at a dose of 1.0 U kg^-1^ body weight. Blood glucose concentrations were measured at 15, 30, 60, and 90 min following insulin injection. All area under curve (AUC) calculations were performed using Prism 10 (Prism 10).

### Indirect calorimetry

Energy expenditure and metabolic parameters were measured using a Comprehensive Lab Animal Monitoring System (CLAMS; Oxymax, Columbus Instruments). Male mice at 15-week of HFD diet (total 20-week old) and age-matched CD controls with no significant weight differences between WT vs. *Enpp1^H362A^* groups were individually housed in metabolic chambers under room temperature and a 12 h light/12 h dark cycle with ad libitum access to food (HFD or CD) and water. Animals were acclimated to the metabolic cages for 24 h prior to data collection. Oxygen consumption (VO₂), carbon dioxide production (VCO₂), respiratory exchange ratio (RER), energy expenditure, locomotor activity, food intake, and water intake were continuously recorded over a 76 h measurement period. Energy expenditure was calculated by the CLAMS software based on VO₂ and VCO₂ measurements. Statistical significance for VO_2_, VCO_2_, food consumption, and energy expenditure was determined by General Linear Model (ANCOVA) with body mass as a covariate. Significance for RER, locomotor and ambulatory movement was determined by two-way ANOVA.

### Glucose uptake assay

#### In vivo

Glucose uptake assay was adapted from a previously published protocol^33^. Briefly, mice fed HFD or age-matched CD controls were fasted for 6 h with free access to water. At time 0, baseline blood glucose levels were measured from tail vein blood using a handheld glucometer (ReliOn Premier). Mice were then administered 2-deoxy-D-[1,2-³H]glucose (³H-2-DG; 100 μCi/kg; PerkinElmer, NET328A001MC) together with human insulin (1 U/kg; Humulin R, Eli Lilly), both dissolved in sterile saline, by intraperitoneal injection. Blood glucose levels were measured again 30 min after injection, after which mice were euthanized. Tissues of interest were rapidly excised, weighed, and stored at −20 °C prior to processing. Samples were homogenized in 300 μL of 1% SDS (Sigma-Aldrich, #436143) for 1–2 min at room temperature using a handheld homogenizer (Fisher Scientific, #15-340-167). Tissue homogenates were transferred to scintillation vials containing 3.5 mL Ultima Gold scintillation fluid, and ³H radioactivity was measured using a scintillation counter with a 1-min counting time per sample. Glucose uptake was quantified as CPM and normalized to tissue weight as indicated.

#### In vitro

Differentiated brown adipocytes were serum-starved for 4 h in Krebs–Ringer HEPES (KRH) buffer composed of 12 mM HEPES (pH 7.5), 120 mM NaCl, 5 mM KCl, 0.33 mM CaCl₂, and 1.2 mM MgSO₄, supplemented with 0.5% fatty acid–free BSA (Sigma-Aldrich, #A4612). Two hours prior to glucose uptake measurements, cGAMP at the indicated concentrations (InvivoGen, #rlrl-nacga23-02, or synthesized in-house) was added. Insulin (100 nM; Abcam, #AB123768) was added to the indicated samples 20 min prior to tracer addition. Glucose uptake was initiated by addition of 500 μM D-glucose (Sigma-Aldrich, #50-99-7) spiked with 2-deoxy-D-[1,2-³H]glucose (³H-2-DG; 1.71 μCi/μL; PerkinElmer, NET328A001MC). After 10 min, cells were washed twice with an ice-cold stop buffer consisting of KRH buffer supplemented with 200 mM D-glucose. Cells were then solubilized in 1% SDS, and lysates were transferred to scintillation vials containing Ultima Gold scintillation fluid. ³H radioactivity was measured using a scintillation counter with a 1-min counting time per sample.

### Histology and lipid staining

Inguinal white adipose tissue (iWAT), brown adipose tissue (BAT), and liver were excised and fixed in neutral buffered formalin (Carolina, #863553) prior to submission to Histo-Tec Laboratory Inc. for paraffin embedding, sectioning, H&E staining. For neutral lipid visualization, liver tissues were embedded in optimal cutting temperature (OCT) compound (Thermo Fischer, #23-730-571) in cryomolds and rapidly frozen on crushed dry ice for 10 min. Frozen sections (10 μm thickness) were prepared using a cryostat at −20 °C and stained with Oil Red O using a commercial kit (Abcam, ab150678) according to the manufacturer’s instructions. Adipocyte size in iWAT and BAT was quantified from H&E-stained sections using ImageJ (v2.0.0)^70^. Lipid droplet area in iWAT was measured by polygonal tracing of 50 randomly selected adipocytes per sample, with three independent samples analyzed per condition. Lipid droplet area in BAT was measured by polygonal tracing of 25 randomly selected unilocular adipocytes per sample, with three independent samples analyzed per condition.

### Circulating insulin, glucagon, and free fatty acid measurement

Blood samples were collected from mice, allowed to clot at room temperature, and centrifuged to obtain serum. Serum insulin and glucagon concentrations were measured using a Mouse Insulin ELISA Kit (Abcam, ab285341) and a Mouse Glucagon ELISA Kit (Fisher Scientific, #50-194-7903), respectively, according to the manufacturers’ instructions. Circulating free fatty acid (FFA) levels were quantified using a fluorometric Free Fatty Acid Assay Kit (Cayman Chemical, #700310). Absorbance or fluorescence signals were measured using a microplate reader, and analyte concentrations were calculated from standard curves generated in parallel.

### Core body temperature measurement

Core body temperature was measured in conscious mice at room temperature using a lubricated rectal temperature probe connected to a digital thermometer (Fischer Scientific, #9713069). Measurements were performed by gently inserting the probe approximately 2 cm into the rectum and recording the stabilized temperature. All measurements were conducted at the same time of day to minimize circadian variation, and mice were allowed to acclimate to the procedure prior to data collection.

### Tissue isolation and immune cell preparation

Brown adipose tissue (BAT) and inguinal white adipose tissue (iWAT) were harvested, mechanically minced, and digested in 5 mL of digestion buffer consisting of RPMI 1640 medium (Thermo Fisher, #C22400500CP) supplemented with 10% heat-inactivated FBS (R&D Systems, #S12450H), collagenase IV (1 mg/mL; Sigma-Aldrich, #C5138), and DNase I (20 μg/mL; Roche, #10104159001). Digestions were performed for 1 h at 37 °C with constant agitation. Cell suspensions were passed through 100-μm sterile cell strainers to remove undigested tissue, and enzymatic digestion was quenched by addition of 5 mL FACS buffer (PBS containing 2% heat-inactivated FBS and 2 mM EDTA). Samples were centrifuged at 500 × g for 5 min at 4 °C, supernatants were discarded, and cell pellets were resuspended in 3 mL of 40% Percoll in PBS and laid below 1 mL of 70% Percoll in RPMI. Density gradient centrifugation was performed at 600 × g for 20 min at room temperature (acceleration 6, deceleration 2). Cells at the 40%–70% Percoll interface were collected, washed in 9 mL FACS buffer, and centrifuged at 500 × g for 5 min at 4 °C. Cells were counted and used for subsequent antibody staining and flow cytometric analysis.

### 2’3’-cGAMP (cGAMP) measurement

For cGAMP measurements in tissue, BAT from HFD-fed mice and age-matched CD controls was excised, snap-frozen in liquid nitrogen, and stored at –80 °C. For analysis, tissues were thawed on ice, and 30–150 mg samples were weighed and placed into impact-resistant screw-cap tubes (USA Scientific, #1420-9600) containing lysis buffer (100 mg/mL) composed of T-PER Tissue Protein Extraction Reagent (Thermo Fisher, #78510) supplemented with ENPP1 inhibitor STF-1084^14^ (100 nM), protease inhibitor cocktail (Roche, #05892970001), and phosphatase inhibitor (Roche, #4906845001). Tissue homogenization was performed using a bead mill homogenizer (Fisher Scientific, #15-340-163) under the following settings: speed 4.5, 30 s, dwell 20 s, 5 cycles. Homogenates were centrifuged at 16,000 × g for 20 min at 4 °C, and the supernatants were collected. cGAMP levels were quantified using a commercially available ELISA kit (Cayman, #501700) according to the manufacturer’s instructions. Lysates were diluted 100 times to ensure measurements fell within the linear range of the assay.

For cGAMP measurements in cells, brown preadipocytes between day 0 and day 2 after seeding prior to differentiation induction, and differentiated brown adipocytes between day 4 and day 6 of differentiation were used. Cells were seeded in 250 μL of differentiation medium containing STF-1623^39^ (1 μM) or PBS as a control. On the day of collection, culture media were harvested and centrifuged at 1,000 × g for 3 min to remove cell debris before supernatant was collected. Remaining cells were washed with PBS and lysed in M-PER Extraction Reagent (Thermo Fisher, #78501) with or without STF-1623 (1 μM). Extracellular and intracellular cGAMP were quantified using a commercial ELISA kit (Cayman, #501700) following the manufacturer’s instructions.

### Cytoplasmic mtDNA quantification

Detection of mitochondrial DNA (mtDNA) was adapted from a previously published protocol^71^. Brown preadipocytes or differentiated adipocytes cultured in 10-cm dishes were trypsinized, washed with PBS, and divided into two fractions. One fraction containing one-fifth of the cells was lysed in 500 μL SDS lysis buffer (20 mM Tris-HCl, pH 8.0, 1% SDS, supplemented with protease inhibitors) and boiled at 95 °C for 15 min to generate whole-cell extracts (WCE). The remaining four-fifths of the cells were resuspended in 600 μL digitonin lysis buffer (50 mM HEPES, pH 7.4, 150 mM NaCl, 20 μg/mL digitonin, supplemented with protease inhibitors) and homogenized using a Dounce homogenizer until approximately 40–50% cell death was confirmed by Trypan Blue staining (Invitrogen, #T10282). Homogenates were centrifuged twice at 600 × g for 5 min to remove nuclei and unbroken cells, and the supernatants were collected. Samples were subsequently centrifuged at 7,000 × g for 10 min, and the resulting supernatants were collected as the cytosolic fraction. DNA from WCE and cytosolic fractions was purified using the DNeasy Blood and Tissue Kit (Qiagen, #69504). Purity of the cytosolic fraction was validated by immunoblotting for the cytosolic marker GAPDH (Cell Signaling Technology, #2118), mitochondrial marker TOM20 (Proteintech, #15793A), and nuclear marker histone H3 (Cell Signaling Technology, #14269). Mitochondrial DNA was quantified by qPCR using primers directed against mouse **Cytb** (F: ATTCCTTCATGTCGGACGAG; R: ACTGAGAAGCCCCCTCAAAT), **D-loop** (F: AATCTACCATCCTCCGTGAAACC; R: TCAGTTTAGCTACCCCCAAGTTTAA), **mt-Tf** (F: CGTTAGGTCAAGGTGTAGCC; R: CCAGACACACTTTCCAGTATG), and **ND4** (F: AACGGATCCACAGCCGTA; R: AGTCCTTCGGGCCATGATT). Nuclear DNA was quantified using primers against **H19** (F: GTCCACGAGACCAATGACTG; R: GTACCCACCTGTCGTCC). Relative cytosolic mtDNA levels were calculated by normalizing absolute mtDNA abundance in the cytosolic fraction to absolute H19 levels measured in the corresponding WCE.

### Cellular viability measurement

Cell viability was assessed using the CellTiter-Glo Luminescent Cell Viability Assay (Promega, #G9241). Cells were incubated with the reagent according to the manufacturer’s instructions, and luminescence was measured using a microplate reader and normalized to the indicated controls.

### Lipolysis assay

Differentiated brown adipocytes were washed twice with Krebs–Henseleit Ringer buffer (KHRB) composed of 30 mM HEPES (pH 7.5), 10 mM NaHCO₃, 4 mM K₂HPO₄, 120 mM NaCl, 1 mM MgSO₄, and 1 mM CaCl₂. Cells were then incubated in KHRB supplemented with 2% fatty acid–free BSA (Sigma-Aldrich, #A4612) in the presence of norepinephrine (Sigma-Aldrich, #A0937) or cGAMP (InvivoGen, #rlrl-nacga23-02, or synthesized in-house^42^) at the indicated concentrations for 2 h. Culture media were collected and heat-inactivated at 65 °C for 10 min. Glycerol release was quantified using a commercial assay kit (Abcam, #133130) according to the manufacturer’s instructions.

### Western blotting

Cell lysates were resolved by SDS–polyacrylamide gel electrophoresis using precast gels (GenScript, #M42015) and transferred onto nitrocellulose membranes using a dry transfer system (Bio-Rad, #1704150). Membranes were blocked and incubated overnight at 4 °C with primary antibodies diluted 1:1,000, including antibodies against tubulin (CST, #2144), STING (CST, #13647), phospho-STING (CST, #72971), insulin receptor β (CST, #3025), phospho-IRβ (CST, #3024), IRS1 (CST, #3194), phospho-IRS1 (CST, #2388), AKT (CST, #2920), phospho-AKT (CST, #4060), GAPDH (CST, #2118), TOM20 (Proteintech, #15793A), histone H3 (CST, #14269), and ENPP1 (Abcam, #240653). Membranes were washed three times in TBS-T (1× TBS containing 0.1% Tween-20) and incubated for 1 h at room temperature with IRDye-conjugated secondary antibodies (IRDye 800CW goat anti-rabbit, LI-COR, #926-32211; IRDye 680RD goat anti-mouse, LI-COR, #926-68070). After three additional washes in TBS-T, immunoreactive bands were visualized using a LI-COR Odyssey infrared imaging system.

### Quantitative real-time PCR (qRT-PCR)

Total RNA was isolated from cells using the Zymo Quick-RNA Miniprep Kit (Zymo Research) according to the manufacturer’s instructions. 1 μg of RNA were reverse-transcribed into cDNA using Maxima Reverse Transcriptase (Thermo Fisher, #EP0725) following the manufacturer’s protocol. Quantitative PCR was performed using TaqMan Gene Expression assays (Applied Biosystems) on a real-time PCR system. Probes from Thermo Fisher include the following: *Ucp1* Mm01244861_m1), *Isg15* (Mm01705338_s1), *Cxcl10* (Mm00445235_m1), *Ifit1* (Mm07295796_m1). Relative gene expression levels were calculated using the ΔΔCt method and normalized to the endogenous control gene *Gapdh* (Thermo Fisher, #Mm9999991_g1)

### Flow cytometry and fluorescence-activated cell sorting (FACS)

After incubation with 5 μg/ml of anti-CD16/32 (2.4G2, BD Biosciences) for 10 min at 4°C, cells were stained with antibody cocktails (around 2×106 cells in 100 μl) for 20 min at 4°C. Dead cells were excluded with Fixable Viability Dye eFluor 780 (Thermo Fisher, Cat# 65-0865-18). Antibodies used to stain immune cells are listed **Table 1** below. Flow cytometry was performed on Attune NxT (Thermo Fisher), and the data were analyzed with FlowJo 10.7.1 software (TreeStar). Cell sorting was performed using BD FACSAria Fusion with a 100-μm nozzle at 4°C.

**Table 1.** Antibody Information.

| **Table 1. Antibody Information** |  |  |  |  |  |
| --- | --- | --- | --- | --- | --- |
| **Antibody** | **Fluorochrome** | **Clone** | **RRID** | **Source** | **Concentration** |
| Anti-CD45 | BV650 | 30-F11 | AB_2565884 | BioLegend | 0.25 µg/mL |
| Anti-CD45 | AF700 | OX-1 | AB_10682900 | BioLegend | 2.5 µg/mL |
| Anti-CD11b | BV510 | M1/70 | AB_2561390 | BioLegend | 1 µg/mL |
| Anti-CD11b | FITC | M1/70 | AB_312788 | BioLegend | 2.5 µg/mL |
| Anti-CD11c | PE | N418 | AB_313776 | BioLegend | 1 µg/mL |
| Anti-F4/80 | APC | QA17A29 | AB_2832548 | BioLegend | 1 µg/mL |
| Anti-Ly-6G/Ly-6C (Gr-1) | BV421 | RB6-8C5 | AB_10900232 | BioLegend | Unspecified, 1:200 dilution |
| Anti-Ly-G6 | BV605 | 1A8 | AB_2565880 | BioLegend | 1 µg/mL |
| Anti-MHC-II | BV510 | M5/114.15.2 | AB_2561397 | BioLegend | 1 µg/mL |
| Anti-CD3 | FITC | 17A2 | AB_312660 | BioLegend | 2.5 µg/mL |
| Anti-NK1.1 | PE | PK136 | AB_313394 | BioLegend | 1 µg/mL |
| Anti-B220 | BV510 | RA3-6B2 | AB_2561394 | BioLegend | 1 µg/mL |

### RNA sequencing (RNA-seq)

For bulk RNA-seq of brown adipocytes with or without cGAMP treatment, cells were lysed in TRIzol reagent (Invivogen, #15596018). Total RNA was isolated from cells using the Quick-RNA Miniprep Kit (Zymo Research) according to the manufacturer’s instructions. RNA samples were submitted to Beijing Genomics Institute (BGI), Inc. for library preparation and high-throughput sequencing. Sample QC was performed with Agilent Tapestation 4200 to identify samples with RIN > 7. 200 ng qualified RNA from each sample was processed for library preparation. In brief, mRNA enrichment was performed on total RNA using oligo(dT)-attached magnetic beads. The enriched mRNA with poly(A) tails was fragmented using a fragmentation buffer, followed by reverse transcription using random N6 primers to synthesize cDNA double strands. The synthesized double stranded DNA was then end-repaired and 5’-phosphorylated, with a protruding ’A’ at the 3’ end forming a blunt end, followed by ligation of a bubble-shaped adapter with a protruding ’T’ at the 3’ end. The ligation products were PCR amplified using specific primers. The PCR products were denatured to single strands, and then single-stranded circular DNA libraries were generated using a bridged primer. The constructed libraries were quality-checked and sequenced after passing the quality control. The library was amplified with phi29 to make DNA nanoball (DNB) which had more than 300 copies of one molecule. The DNBs were loaded into the patterned nanoarray and pair end 150 bases reads were generated and sequenced by synthesis. The sequencer used is MGI G400 platform. The sequencing data, also known as raw reads or raw data, were subjected to quality control (QC) to determine whether the sequencing data were suitable for subsequent analysis. After QC, the clean reads were aligned to the reference sequences. After alignment, the alignment results were subjected to a second QC to evaluate the alignment quality, including alignment rate and distribution of reads on the reference sequences. After that, gene quantification analysis, various analyses based on gene expression levels (such as principal component analysis, correlation analysis, differential gene screening, etc.) were performed. Differential expressed genes between samples were further analyzed for gene ontology (GO) functional enrichment, pathway enrichment, clustering, protein interaction networks, and transcription factors. HISAT (Hierarchical Indexing for Spliced Alignment of Transcripts) is a software used for aligning RNA-seq reads to a reference genome. Clean data was aligned to the reference gene set using Bowtie2 (v2.3.4.3). Gene expression quantification was performed using RSEM (v1.3.1) software, and gene expression clustering heatmaps across different samples were generated using heatmap (v1.0.8). DESeq2 (v1.4.5) (or DEGseq or PoissonDis) was employed for differential gene detection, with criteria set as Q value ≤ 0.05 or FDR ≤ 0.001. Sequencing data were visualized and analyzed using the OEBiotech cloud platform (cloud.oebiotech.com).

For bulk RNA sequencing of brown adipose tissue macrophages, 1,000 fluorescence-activated cell sorting (FACS)–purified macrophages were lysed in 10 μL of TCL buffer (Qiagen, #1031576) supplemented with 1% 2-mercaptoethanol (Sigma-Aldrich, #M3148-25ML). Whole-transcriptome amplification (WTA) was performed using a published SMART-Seq2 protocol^72^. Amplified cDNA was purified and quality-controlled using a Bioanalyzer (Agilent). Five biological replicates per genotype were submitted to Genewiz, Inc. for library preparation and high-throughput sequencing. The DNA was quantified using the Qubit 2.0 Fluorometer (ThermoFisher Scientific, Waltham, MA, USA). The NEBNext Ultra II DNA Library Prep Kit for Illumina (New England Biolabs, Ipswich, MA, USA), including clustering and sequencing reagents, was utilized according to the manufacturer’s recommendations. Briefly, adapters were ligated after adenylation of the 3’ ends followed by enrichment by limited cycle PCR. DNA libraries were validated using the Agilent TapeStation (Agilent Technologies, Palo Alto, CA, USA) and were quantified using Qubit 2.0 Fluorometer. The DNA libraries were also quantified by real time PCR (KAPA Biosystems, Wilmington, MA, USA). The sequencing library was clustered onto a flowcell. After clustering, the flowcell was loaded onto the Illumina NovaSeq XPlus instrument according to the manufacturer’s instructions. The samples were sequenced using a 2x150bp Paired End (PE) configuration. Image analysis and base calling were conducted by the NovaSeq Control Software (NCS). Raw sequence data (.bcl files) generated from Illumina NovaSeq was converted into fastq files and de-multiplexed using Illumina bcl2fastq 2.20 software. One mis-match was allowed for index sequence identification. After investigating the quality of the raw data, sequence read will be trimmed to remove possible adapter sequences and nucleotides with poor quality using Trimmomatic v.0.36. The trimmed reads will be mapped to the GRCm38 reference genome available on ENSEMBL using the STAR aligner v.2.5.2b. The STAR aligner is a splice aligner that detects splice junctions and incorporates them to help align the entire read sequences. BAM files will be generated as a result of this step. Unique gene hit counts will be calculated by using feature Counts from the Subread package v.1.5.2. Only unique reads that fall within exon regions will be counted. After extraction of gene hit counts, the gene hit counts table was used for downstream differential expression analysis. Using DESeq2, a comparison of gene expression between the groups of samples was performed. The Wald test was used to generate p-values and Log2 fold changes. Genes with adjusted p values < 0.05 and absolute log2 fold changes > 1 were called as differentially expressed genes for each comparison. A gene ontology analysis was performed on the statistically significant set of genes by implementing the software GeneSCF. The goa_human GO list was used to cluster the set of genes based on their biological process and determine their statistical significance. A PCA analysis was performed using the “plotPCA” function within the DESeq2 R package. The plot shows the samples in a 2D plane spanned by their first two principal components. The top 500 genes, selected by highest row variance, were used to generate the plot. Sequencing data were visualized and analyzed using the OEBiotech cloud platform (cloud.oebiotech.com).

### hENPP1 Purification and hIR Interaction Analysis

Purification of His-tagged hENPP1 (WT and K173Q) were expressed in HEK293 cells and purified via nickel-affinity chromatography. For pulldown assays, purified WT-hENPP1-His or K173Q-hENPP1-His were incubated with hIR-Tsi-Flag in the presence or absence of insulin. In parallel, co-purification analysis was performed using HEK293 cells co-expressing K173Q-hENPP1-His and hIR-Tsi-Flag. For size exclusion chromatography, a mixture of purified K173Q-hENPP1 and hIR was analyzed using a Superose 6 column.

### TurboID proximity labeling

The experiment was adapted from a previous publication^73^. Briefly, 293T *ENPP1^-/-^* cells^14^ were transiently transfected with MCS-13X Linker-BioID2-HA (empty vector, WT, or K173Q; Addgene 74224) using FuGENE 6 (Promega, #E2691) and Opti-MEM (Thermo Fischer, #31985088). At 24 h post-transfection, the culture media was supplemented with 50 µM biotin and incubated for an additional 24 h to induce proximity-dependent biotinylation of neighboring proteins. Cells were washed three times with ice-cold PBS and lysed in RIPA buffer (Fisher Scientific, #PI89901) supplemented with a protease inhibitor cocktail (Thermo Fischer, #78425). Lysates were incubated with streptavidin-conjugated beads at 4 °C overnight with gentle rotation. Following incubation, the beads were washed three times with PBS. The bead-bound proteins were submitted to the Beijing Genomics Institute (BGI), Inc. for on-bead digestion followed by liquid chromatography–tandem mass spectrometry (LC–MS/MS) analysis.

### ENPP1 cGAMP activity assay by thin-layer chromatography assay

293T *ENPP1^-/-^* cells^42^ were transiently transfected with pcDNA-hENPP1-FLAG (WT or K173Q) using FuGENE 6 (Promega, #E2691) and Opti-MEM (Thermo Fischer, #31985088). 24-48 hours post-transfection, cells were harvested and lysed in 100 μL of lysis buffer (10 mM Tris pH 9.0, 150 mM NaCl, 10 μM ZnCl_2_, 1% NP-40). For western blot analysis, 30 μL aliquots are mixed with 5x reducing sample buffers, heated at 95 °C for 5 min, and sonicated. To ensure comparable enzymatic loading, various lysate volumes were screened via western blot; volumes yielding equivalent WT and K173Q expression levels were selected for subsequent activity assayed. Normalized lysate samples were incubated with 1 μM cGAMP (spiked with trace [^32^P] cGAMP) in ENPP1 activity buffer (50 mM Tris pH 9, 250 mM NaCl, 0.5 mM CaCl_2_, 1 μM ZnCl_2_). At the indicated time points, 1 μl aliquots of the reaction were quenched by spotting onto HP-TLC silica gel plates (Millipore, Cat# 1.05548.0001). Plates were developed in a mobile phase consisting of 85% ethanol and 5 mM NH_4_HCO_3_), then exposed to a phosphor screen (GEBAS-IP MS). The screens were imaged using a Typhoon 9400 scanner, and the ^32^P signal was quantified using ImageJ.

### Statistical information

Prism 10 (GraphPad Software) was used to perform two-tailed *t* tests and two-way mixed model ANOVA test. Indirect calorimetry significance was calculated using CalR (version 1.3)^74^: Statistical significance for VO_2_, VCO_2_, food consumption, and energy expenditure was determined by General Linear Model (ANCOVA) with body mass as a covariate. Significance for RER, locomotor and ambulatory movement was determined by two-way ANOVA.Graphs show means and standard deviation (± s.d.) or standard error of the mean (± s.e.m). In cGAMP degradation assays and pharmacokinetics analyses, half-life was obtained by one phase exponential decay fitting with Prism software. Statistical significance, group size, and experimental details are described in the figure legends.

### Generative AI

During the preparation of this work the author(s) used Gemni in order to proofread the writing. After using this tool/service, the author(s) reviewed and edited the content as needed and take(s) full responsibility for the content of the published article.

## Data Availability

Raw and processed data for RNA sequencing in Figure 5 can be accessed via GSE315247. Raw and processed data for RNA sequencing in Figure 6 can be accessed via GSE315384. All data reported in this paper will be shared by the lead contact upon request. Any additional information required to reanalyze the data reported in this paper is available from the lead contact upon request.

## Acknowledgements

We thank Li Lab members, A. Pawluk, B. Plosky, and the Scientific Publication Team at Arc Institute for feedback throughout the course of this study. We thank our funding support, including Arc Institute (LL, SW, YG, YL, GG, GS, VS, ML), Knight-Henessey Scholarship (ML). CLAMS Oxymax Metabolic Cages Data was obtained using the services of the Stanford Islet Research Core facility of the Stanford Diabetes Research Center which is supported by the National Institute of Diabetes and Digestive and Kidney Diseases of the National Institutes of Health under Award Number P30KD116074.

## Author Contributions

Conceptualization: SW, LL; Methodology: SW, YG, YL, VS, KS, LL; Materials: GG, GS; Investigation: SW, YG, ML; Visualization: SW, LL; Funding acquisition: ML, KJS, LL; Supervision: KJS, LL; Writing – original draft: SW, LL; Writing – review & editing: SW, LL

## Competing Interests

L.L. and S.W. have filed one patent application on methods of use of ENPP1 inhibition (PCT/US2024/024497). L.L. is an inventor of two ENPP1 inhibitors patents (PCT/US2020/015968 and PCT/US2018/050018) that were licensed to Angarus Therapeutics. Angarus Therapeutics’ assets were purchased by Cyana Therapeutics.

## Materials & Correspondence

Further information and requests for resources and reagents should be directed to and will be fulfilled by the lead contact, Lingyin Li. Unique reagents generated in this study are available from the lead contact with a completed Materials Transfer Agreement.

---

**Figure S1.**
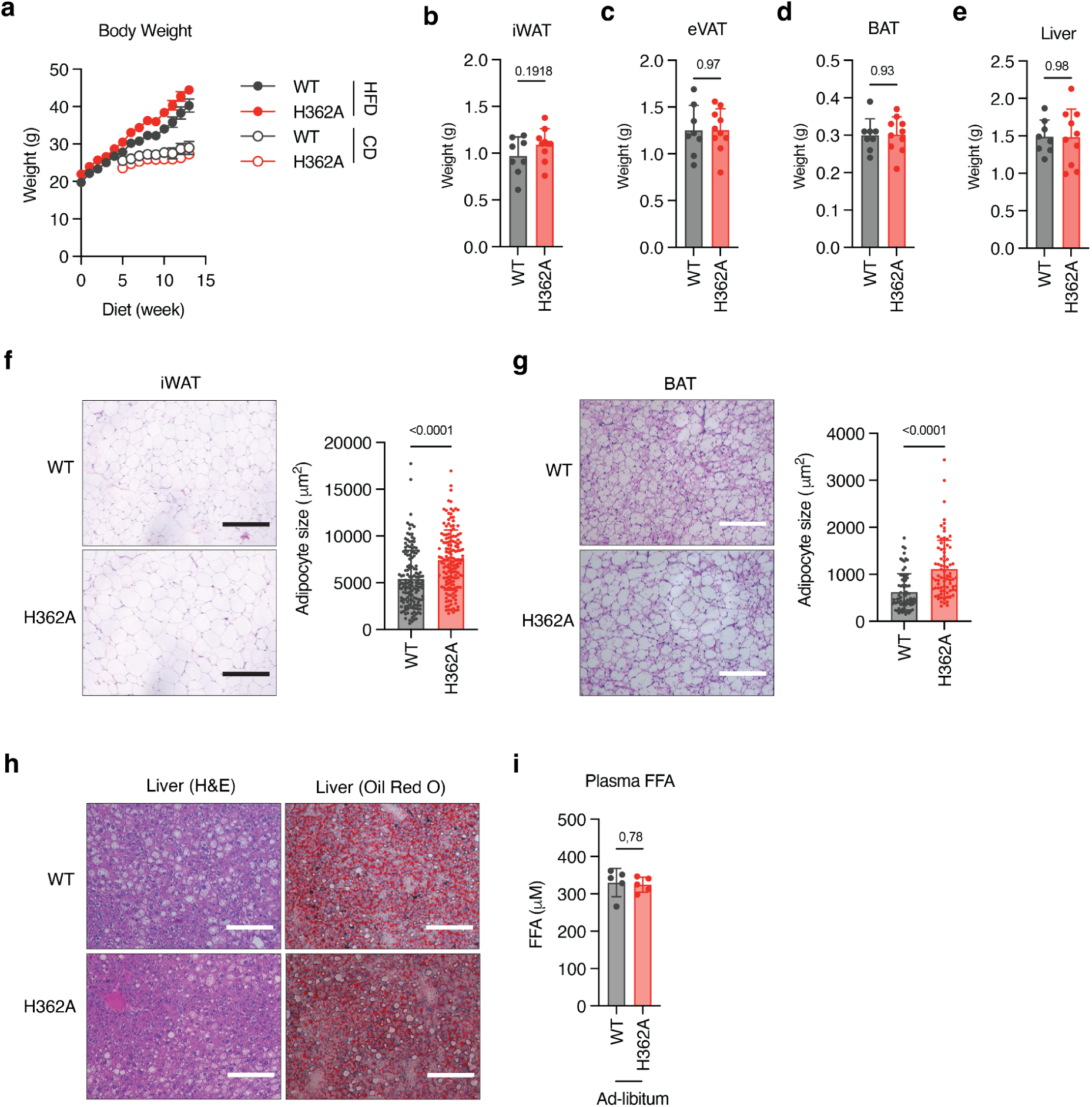
*(Related to Fig. 2)* **a**, Absolute change in body weight of male mice fed a CD or HFD for 14 weeks and housed at room temperature. *n* = 9–10 mice for HFD and *n* = 5 mice for CD. Data are presented as mean ± s.e.m. **b–e**, Weights of iWAT, eVAT, BAT, and liver from male mice after 14 weeks of HFD feeding and housed at room temperature. *n* = 8 or 10 mice per group. Data are presented as mean ± s.d. **f**, Representative H&E staining of iWAT from male mice after 14 weeks of HFD feeding (left) and quantification of adipocyte size (right). Adipocyte area was determined by manual tracing of 50 randomly selected lipid droplets per mouse; three mice per genotype were analyzed. Scale bar, 300 μm. Data are presented as mean ± s.d. **g**, Representative H&E staining BAT from male mice after 14 weeks of HFD feeding (left) and quantification of unilocular adipocyte size (right). Adipocyte area was determined by manual tracing of 25 randomly selected lipid droplets within unilocular brown adipocytes per mouse; three mice per genotype were analyzed. Scale bar, 100 μm. Data are presented as mean ± s.d. **h**, Representative H&E and Oil Red O staining of liver sections from male mice after 14 weeks (H&E) or 9 weeks (Oil Red O) of HFD feeding. Scale bar, 150 μm. **i**, Non-fasted plasma FFA concentrations in male mice after 9 weeks of HFD feeding and housed at room temperature. *n* = 5 mice. Data are presented as mean ± s.d. Statistical significance was assessed using two-sided unpaired *t* tests. H&E: haematoxylin and eosin..

**Figure S2.**
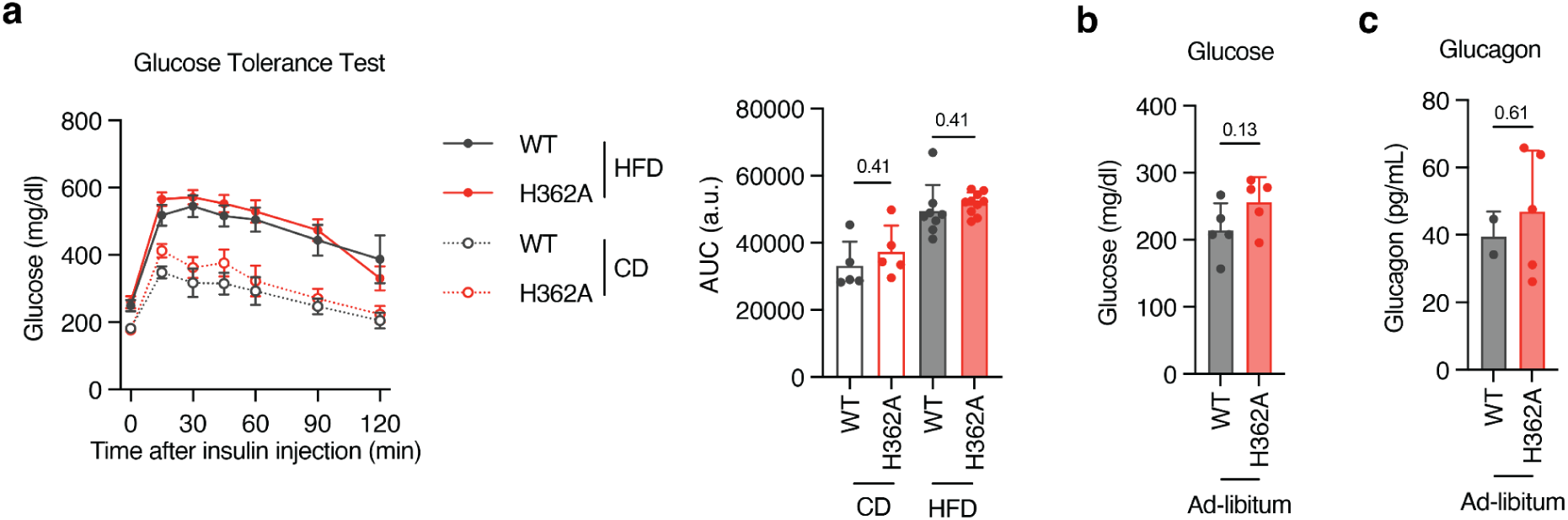
*(Related to Fig. 3)* **a**, Glucose tolerance tests (GTTs) performed after 8 or 13 weeks of HFD or CD feeding in male mice housed at room temperature. Mice were fasted for 5–6 h prior to intraperitoneal injection of glucose (1.5 g kg⁻¹). Blood glucose concentrations were measured over time and AUC was calculated. *n* = 4–5 mice for CD and *n* = 8–9 mice for HFD. Line graphs show mean ± s.e.m.; bar graphs show mean ± s.d. **b**, Non-fasted blood glucose concentrations in male mice after 8 weeks of HFD feeding and housed at room temperature. *n* = 4–5 mice. Data are presented as mean ± s.d. **c**, Non-fasted serum glucagon concentrations in male mice after 8 weeks of HFD feeding and housed at room temperature. *n* = 2–5 mice. Data are presented as mean ± s.d. Statistical significance was assessed using two-sided unpaired ***t*** tests. WT, widetype; H362A, *Enpp1^H362A^*; CD, chow diet; HFD, high-fat diet; GTT, glucose tolerance test.

**Figure S3.**
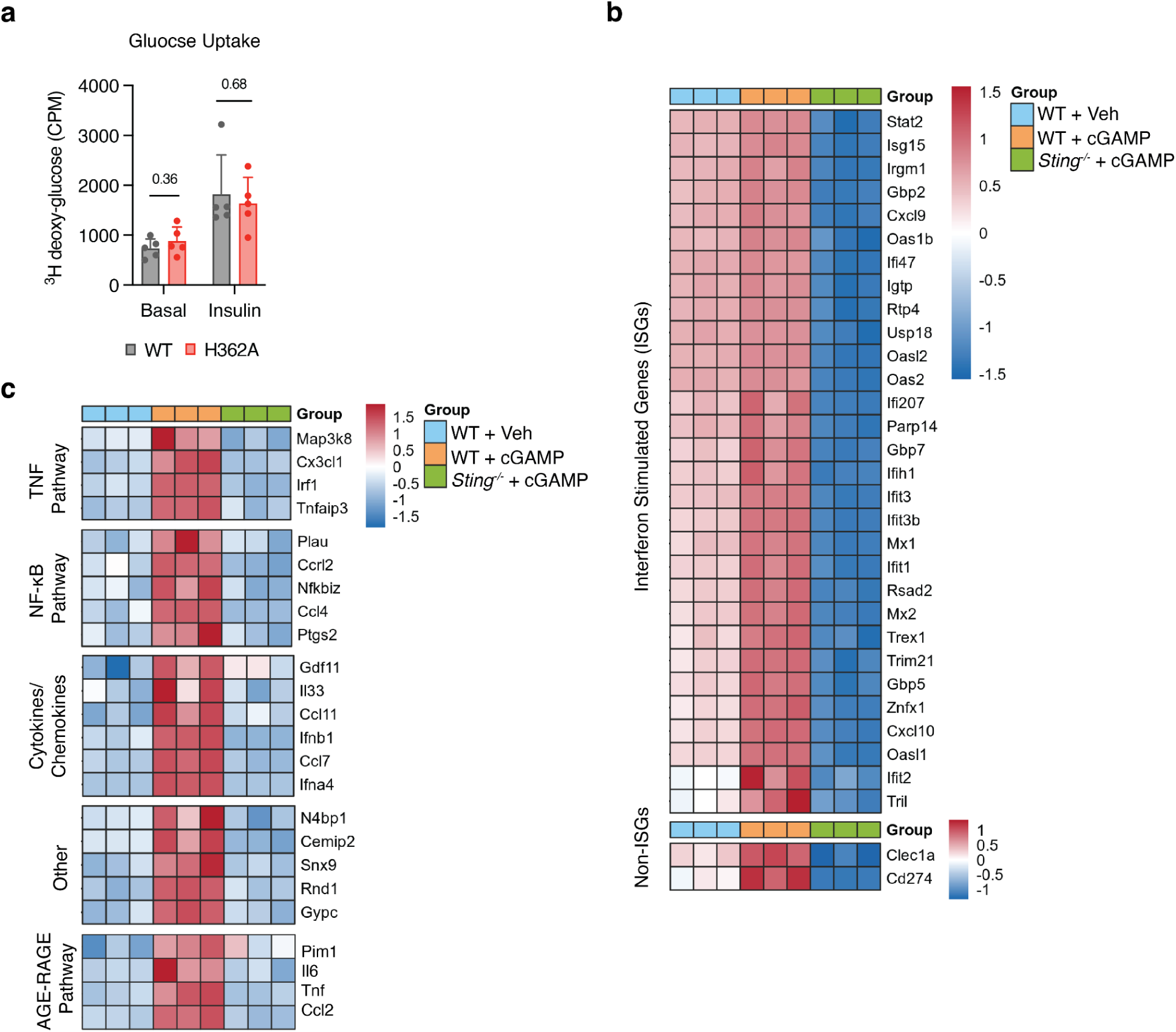
*(Related to Fig. 5)* **a**, ^3^H-2-DG uptake in WT or H362A brown adipocytes under basal conditions or after 20 min insulin (100 nM) treatment. *n* = 5 independent experiments. Data are presented as mean ± s.d. **b,c**, Bulk RNA sequencing of WT brown adipocytes treated with vehicle (WT + Veh), WT brown adipocytes treated with cGAMP (WT + cGAMP), and *Sting^-/-^* brown adipocytes treated with cGAMP (*Sting^-/-^* + cGAMP). Cells were treated with 50 µM cGAMP for 2 h and, where indicated, and 100 nM insulin for 30 min. *n* = 3 biological replicates per group. Sequencing was performed as pair-ended reads on a DNBseq-G400 platform. Expression was normalized to transcripts per million (TPM). Significantly altered genes were defined as Q < 0.05. **b**, Heat map of genes significantly altered between WT + Veh and WT + cGAMP, that are basally expressed in WT + Veh compared with *Sting^-/-^* + cGAMP, including mostly ISGs. Color scale represents z-scores. **c**, Heat map of genes significantly altered between WT + Veh and WT + cGAMP, that are not basally expressed in WT + Veh compared with *Sting^-/-^* + cGAMP, including genes in TNF pathway, NF-κB pathway, cytokines and chemokines, and AGE-RAGE pathway. Color scale represents z-scores Statistical significance was assessed using two-sided unpaired *t* tests comparing WT and *Enpp1^H362A^* mice. ^3^H-2-DG: 2-deoxy-D-[1,2-^3^H]glucose; H362A, *Enpp1^H362A^;* ISG, interferon-stimulated genes; TNF, tumor necrosis factor; NF-κB, nuclear factor kappa-light-chain-enhancer of activated B cells.

**Figure S4.**
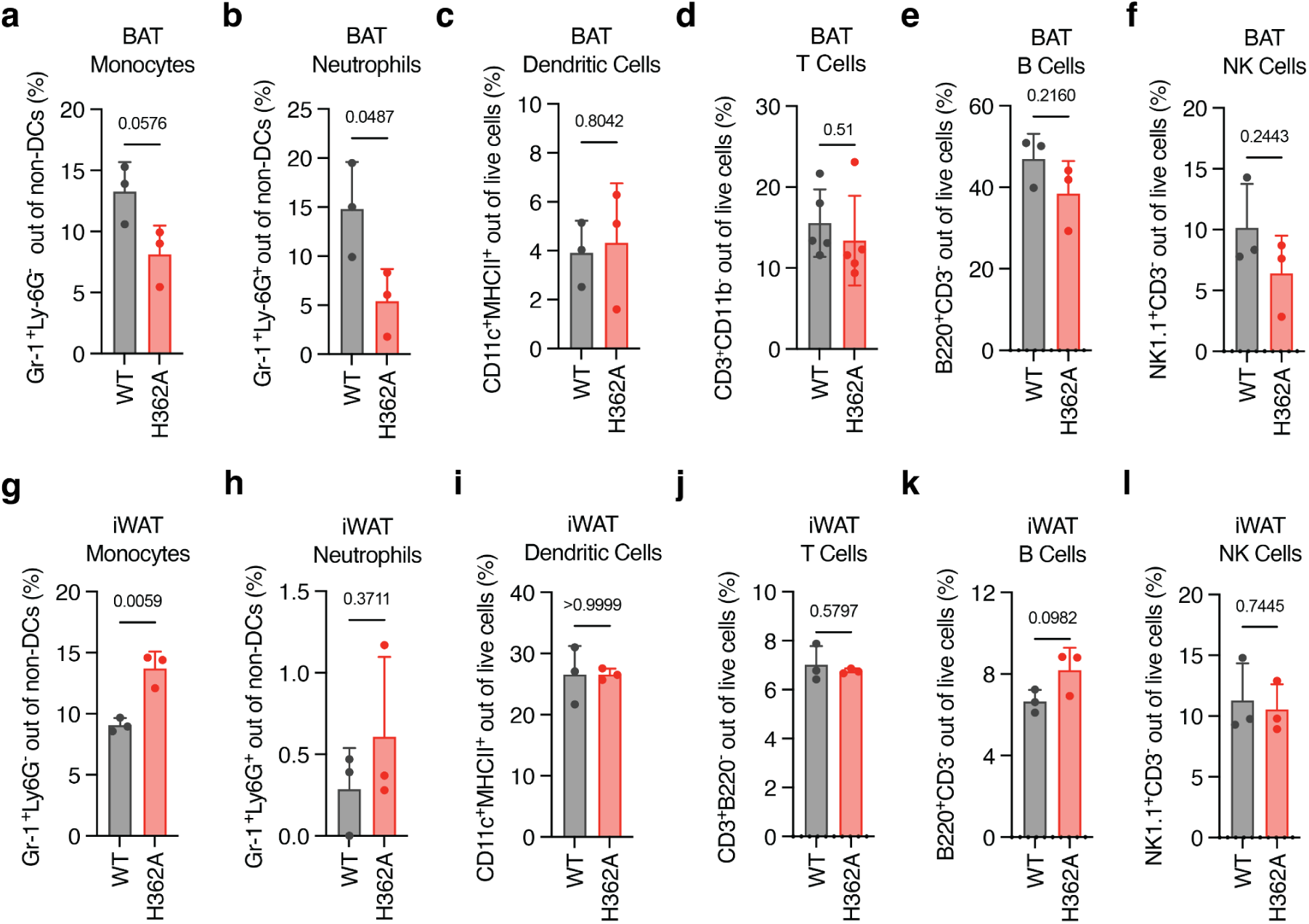
*(Related to Fig. 6)* **a**, Percentage of monocytes (Gr-1⁺Ly-6G⁻) among non-DCs in BAT from male WT and H362A mice housed at room temperature and fed an HFD for 15 weeks. *n* = 3 mice per group. Data are presented as mean ± s.d. **b**, Percentage of neutrophils (Gr1⁺Ly6G⁺) among non-DCs in BAT from the same mice in **a**. **c**, Percentage of DCs (CD11c⁺MHCII⁺) among live cells in BAT from the same mice in **a**. **d**, Percentage of T cells (CD3⁺CD11b⁻) among live cells in BAT from male WT and H362A mice housed at room temperature and fed an HFD for 9 weeks. *n* = 5 mice per group. Data are presented as mean ± s.d. **e**, Percentage of B cells (CD3⁻B220⁺) among live cells in BAT from the same mice in **a**. **f**, Percentage of NK cells (CD3⁻NK1.1⁺) among live cells in BAT from the same mice in **a**. **g**, Percentage of monocytes (Gr-1⁺Ly-6G⁻) among non-DCs in iWAT from the same mice in **a**. **g**, Percentage of neutrophils (Gr1⁺Ly6G⁺) among non-DCs in iWAT from the same mice in **a**. **i**, Percentage of DCs (CD11c⁺MHCII⁺) among live cells in iWAT from the same mice in **a**. **j**, Percentage of T cells (CD3⁺CD11b⁻) among live cells in iWAT from the same mice in **a**. **k**, Percentage of B cells (CD3⁻B220⁺) among live cells in iWAT from the same mice in **a**. **l**, Percentage of NK cells (CD3⁻NK1.1⁺) among live cells in iWAT from the same mice in **a**. Statistical significance was assessed using two-sided unpaired ***t*** tests. BAT, brown adipose tissue; DC, dendritic cells; WT, wild type; H362A, *Enpp1^H362A^*; NK, natural killer; iWAT, inguinal white adipose tissue

